# Regulatory logic of human cortex evolution by combinatorial perturbations

**DOI:** 10.1101/2025.04.28.651083

**Authors:** Adrianos Skaros, Alessandro Vitriolo, Oliviero Leonardi, Veronica Finazzi, Marlene F. Pereira, Filippo Prazzoli, Sebastiano Trattaro, Juan Moriano, Daniele Capocefalo, Carlo Emanuele Villa, Michael Boettcher, Cedric Boeckx, Giuseppe Testa

## Abstract

Comparative genomic studies between contemporary and extinct hominins revealed key evolutionary modifications, but their number has hampered a system level investigation of their combined roles in scaffolding modern traits. Through multi-layered integration we selected 15 genes carrying nearly fixed *sapiens*-specific protein-coding mutations and developed a scalable design of combinatorial CRISPR-Cas9 bidirectional perturbations to uncover their regulatory hierarchy in cortical brain organoids. Interrogating the effects of overexpression and downregulation for all gene pairs in all possible combinations, we defined their impact on transcription and differentiation and reconstructed their regulatory architecture. We uncovered marked cell type-specific effects, including the promotion of alternative fates and the emergence of interneuron populations, alongside a core subnetwork comprising *KIF15*, *NOVA1*, *RB1CC1* and *SPAG5* acting as central regulator across cortical cell types.

## Main Text

The availability of high-coverage genomes from our closest extinct relatives (*1–4*) offers the unprecedented opportunity to probe at high resolution the biological foundations underlying the distinctively modern human condition (*5–7*). Among such differences we expect to find changes impacting early brain development, as the fossil record points to a distinct brain ontogeny reflected in the distinctive contemporary *sapiens* globular neurocranium (*8*). As is to be expected from a highly polygenic trait, bioinformatic analyses have provided a long list of plausible variants of interest (*9*), but the precise impact of many of these still await functional validation. Brain organoids appear to be ideally-suited experimental models to capture the nature of early brain growth differences, as they recapitulate fetal brain development with a high degree of fidelity (*10*). Indeed, attempts have been made to interrogate the impact of derived single nucleotide variants by growing brain organoids containing the CRISPR-engineered ancestral variant of interest and comparing them with typically developing human brain organoids (*11–13*).

Until now studies have mostly focused on understanding the impact of *sapiens*-derived alleles in protein-coding regions that are virtually fixed across modern populations compared to the genomes of extinct hominins. Such work has offered valuable insights, but remains limited in several respects. First, current gene-editing methods impose severe limitations regarding the number of ‘ancestralized’ variants that can be studied at once. Yet we know that most biological processes are driven by combinatorial factors. It is therefore imperative to prioritize multiplex experiments in order to approximate the relevant mechanisms. This is especially pressing since many genes harboring distinctive mutations remain poorly studied, depriving us of valuable knowledge concerning the scope of their actions: we want to understand the extent to which the potentially deleterious effect of a mutation in one gene may be buffered by another mutation in another, interacting gene. Second, while variants in protein-coding regions are reasonable initial targets because of the relatively low number of such changes (virtually) fixed across modern human populations, it is becoming increasingly clear that gene regulation (variants in non-coding regions) is a major source of variation across closely related species (*14–16*), and that effects of genetic variation on complex traits act primarily via changes in gene regulation. Finally, in light of our increased grasp of the expanding magnitude of modern human genetic variation and the evidence of multiple bi-directional gene flow among closely related hominin lineages (Fig. 1A), we deem it important to consider not only fixed mutations but also segregating mutations at high frequency (>90% fixed across modern populations) as relevant to capture the “human condition” (*7,9*).

**Figure 1.**
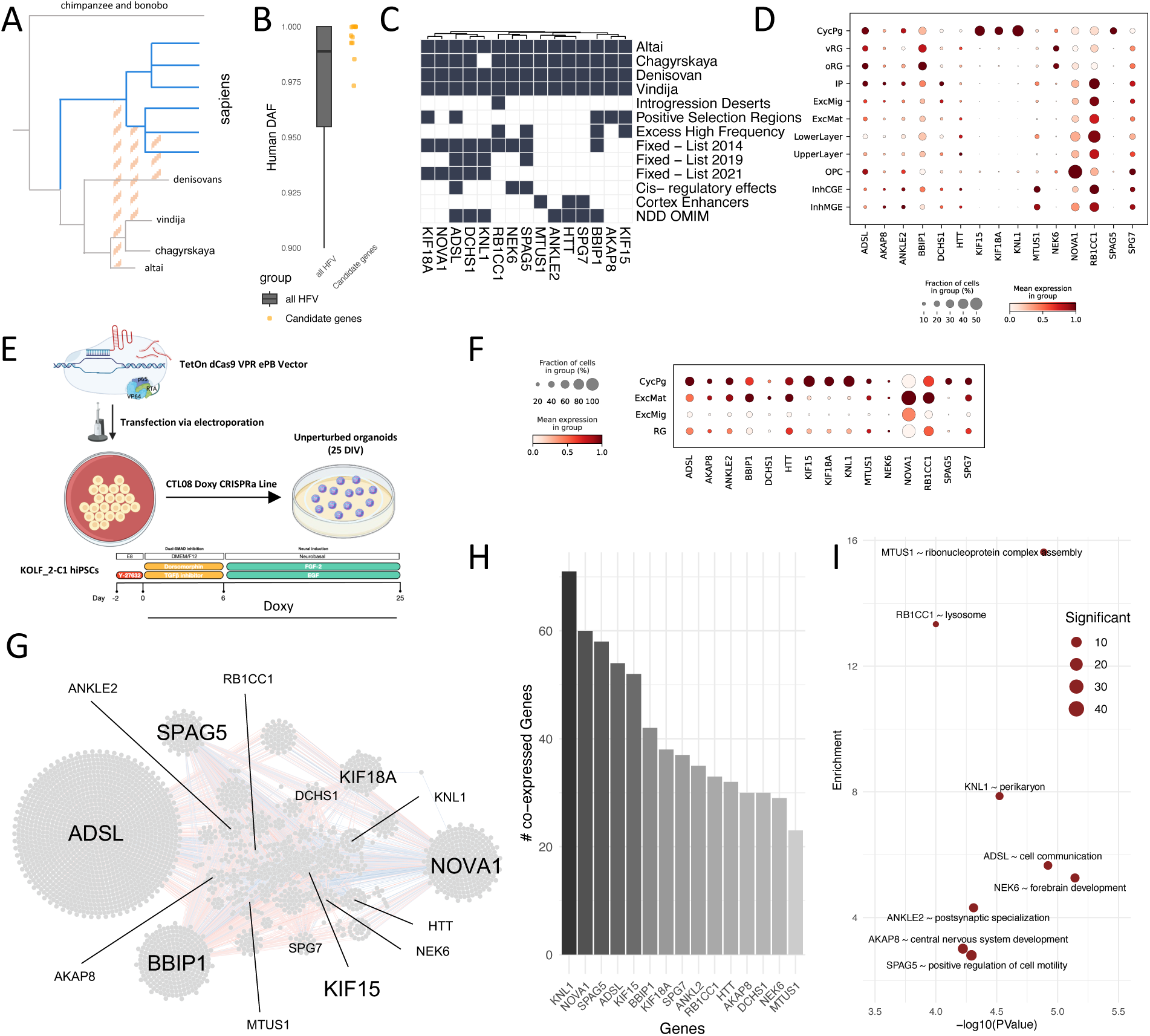
Gene selection and co-expression analysis. A) Scheme of *Homo* evolution and gene flows. Dashed orange lines represent gene flows, blue sticks represent ramifications of *Homo sapiens* populations; B) Boxplot of the distribution frequencies of high-frequency variants (HFV) in grey, next to the distribution (orange dots) of our candidate genes; C) Matrix plot summarizing the features (y-axis) used for gene (x-axis) selection. The first four rows report the presence of variant in Altai, Chagyraskaya, Denisovan, and Vindija genomes. Subsequent rows denote HFV in introgression deserts, signals of positive selection, and excess HF regions, while additional rows highlight fixed mutations identified in ^2,9,11^. The cis-regulatory effect indicates promoter-proximal regulatory variants, “Cortex Enhancers” marks mutations overlapping PsychENCODE cortical enhancers, and “NDD OMIM” identifies genes linked to developmental disorders in Online Mendelian Inheritance in Man. D) Dotplot of average expression per cell type in Polioudakis et al.^58^ single-cell atlas, with nomenclature being largely derived from the original paper. Excitatory neurons (ExcN) were divided into Lower and Upper Layer neurons based on TBR1 expression (see Methods). CycPg = Cycling progenitors, vRG = ventral Radial Glia, oRG = outer Radial Glia, IP = Intermediate Progenitors, ExcMig = excitatory migratory neurons, ExcMat = excitatory maturing neurons, OPC = Oligodendrocyte Precursor Cells, InhCGE and InhMGE = Inhibitory neurons from Caudal and Medial Ganglionic Eminence respectively. G) Co-expression network of core genes, where nodes represent genes and edges denote significant co-expression (see Methods); H) Barplot of the number of uniquely co-expressed genes per gene, based on the top 100 co-expressed genes of each of the 15 candidates; I) Scatterplot of the top GO enrichment derived from the top 100 co-expressed genes of each of our candidates.

For all these reasons we developed a high-throughput approach that aims to illuminate the nature of multiple genes of evolutionary relevance simultaneously, their interactions, and the regulatory logic in which they are embedded. The approach repurposes recently developed orthogonal CRISPR screening approach (*17,18*), based on co-expression of two different Cas9 systems, namely a Streptococcus pyogenes (Sp) based CRISPRa system and a Staphylococcus aureaus Cas9 nuclease. By combining in the same cell Sp Cas9 for gene activation and Sa Cas9 for gene deletion, we performed a dual-screen approach that enables the multiplexed simultaneous overexpression of one gene and deletion of a second gene within the same cell. By using two separate Cas9 enzymes, we ensure that the PAM sequences located at the target site and sgRNA TRACR structure of each sgRNA are distinct, preventing competition between them. Additionally, by employing different promoters for each sgRNA, we avoid erroneous recombination during plasmid propagation in E.coli. By pooling all possible perturbation pairs, this approach enables the reconstruction of directional regulations, as single genes will be perturbed in either direction, first in combination with non-target control sgRNAs, and then in pairs. Our approach breaks new ground in two ways: first, it implements these experimental pipelines in brain organoids, the model that is most ideally suited to probe the nature of the neurobiological differences we are interested in. Second, while our constructs are designed to work with currently commercially available scRNAseq library preparation kits, to achieve the highest guide detection rates we had to bypass standard softwares (see Methods).

### Core candidate mediators of Sapiens cortex evolution

We started with a multilayered analysis to select core genes involved in the evolution of the human cortex, aiming at prioritizing 15 top ranking candidates to obtain the manageable range of cell numbers needed to score, in our multiplexed perturbation design, transcriptome-level phenotypes for the main brain organoid subpopulations. From our catalog of single nucleotide changes distinguishing contemporary humans from other hominins (*9*), we integrated the following subsets and criteria:

i) missense mutations “nearly fixed” in modern human populations, i.e. whereby a threshold of 90% frequency across human populations was set for the derived (i.e. *sapiens*) allele (*9*). In contrast to previous work on protein-coding mutations segregating among hominins (*11,12*), we did not impose the requirement of full fixation because, in line with our original proposition (*9*), recurrent bidirectional gene flow between hominin populations and a more comprehensive assessment of variation across contemporary and extinct humans affect allele frequency distribution^7^, eroding the status of full fixation with a 50% drop in “fully fixed” variants over just a few years (compare *5* and *9*).
ii) single nucleotide changes falling within regulatory regions and expected to impact gene expression at early stages of cortical development, with a higher score for changes found in so-called “regulatory islands” (homolog regions in the Neanderthal/Denisovan genomes in which species-specific, derived variants are not found) (*19,20*), or for which experimental evidence points to gene expression differences in neural progenitors associated with derived variants (*14*).
iii) expression at early stages of cortical development, both from fetal brain data (*21*) and our in-house cortical organoid scRNAseq datasets (*22,23*). This aimed both at a functional enrichment, given that genes with derived variants at a frequency greater than 99.9% in present-day humans are preferentially expressed in the ventricular zone, relative when compared to other hominins (*9* refining an earlier result of *2*) and at the reliable detection of its changes upon perturbations.
iv) evidence of association with neurodevelopmental processes in human populations, including phenotypic associations in the Online Mendelian Inheritance in Man (OMIM) database (https://www.omim.org/).
v) association with signals of positive selection (*24*), presence within large deserts of introgression (regions of the modern human genomes depleted of alleles introgressed from other hominins) (*25,26*), and excess of high frequency derived mutations relative to Neanderthals and Denisovans, controlling for gene length (*9*) (Fig. 1C).

Together, this led us to prioritize 15 genes, with relevant biological properties (see *9* for references and additional discussion) i) ADSL is an enzyme that affects metabolic processes implicated in brain development; ii) AKAP8 and DCHS1 are membrane proteins that contribute to cytoskeletal organization and neuronal migration; iii) ANKLE2 is an inner nuclear membrane ankyrin repeat protein and KNL1 is a phosphatase, both microcephaly associated genes; iv) BBIP1 is a member of the BBSome complex involved in ciliary transport and signaling iv) KIF18A and KIF15, constituents of the kinetochore, kinase NEK6, and and the spindle protein SPAG5, all coordinating cell cycle dynamics; vi) HTT, beyond its role in Huntington’s disease, has been linked to cortical morphogenesis; vii) NOVA1 regulates alternative splicing of neuronal genes; viii) the kinase RB1CC1, the mitochondrial protease SPG7 and microtubule scaffold protein MTUS1 modulate autophagy, mitochondrial and signaling pathways, respectively.

These genes harbor a total of 19 nearly fixed derived missense mutations (3 for *SPAG5*, 2 for *KNL1* and *KIF15*, and 1 for all the other genes), with most of them currently virtually fixed (>99% frequency across contemporary populations) (*11,27*) (Fig.1B).

These core targets span a range of expression patterns along corticogenesis with, at one extreme, three of them (*KIF15*, *KIF18A* and *KNL1*) highly specific for cycling progenitors and at the other extreme *SPG7*, largely depleted in the same progenitors, and a more generalized expression for the other genes (Fig. 1D).

### Line generation and Cortical organoid differentiation

To achieve, in the same cell, simultaneous inactivation of one gene and overexpression of another in all possible combinations, we devised a system based on two separately inducible Cas9 proteins. To separately validate the proficiency of each perturbation axis, we first engineered the standard KOLF2C1 pluripotent stem cell line to stably express the doxycycline-inducible *sp* dCas9-VPR transgene (CRISPRa system) (see Methods) for efficient gene activation upon sgRNAs transduction, and validated the preservation of transgene induction upon extended culture and differentiation into cortical brain organoids (Fig. 1E). We confirmed the inducible overexpression proficiency by targeting the transmembrane ABCG2 (Supp. Fig.1A-B).

Organoids were cultured through 25 days in vitro (DIV), a stage that we and others benchmarked to capture radial glia and early neuronal populations (*10,28-30*). We profiled unperturbed organoids (ie. harboring only the CRISPRa transgene) by single-cell RNA-seq (scRNAseq) to confirm their cell type composition and gather functional information on each gene, by evaluating their co-expression patterns. We identified four major cell populations expressing markers of cycling neural progenitors (CycPg), radial glia (RG), migrating cortical neurons (ExcMig) and maturing excitatory neurons (MatExc) (Fig. S1C-D) and expressing our candidate genes consistently with our observation from fetal datasets (Fig. 1F). We next characterized the activity of our candidate genes, grouping them by cell types, and performing co-expression analysis on pseudobulks (see Methods). While they behaved differently in terms of sheer number of co-expressed genes, our targets shared enough of them to allow prediction of a core co-expression network (Fig. 1G, Fig.S1E), with *KNL1*, *NOVA1 SPAG5*, *ADSL* and *KIF15* ranking at the top with more than 50 co-expressed genes each, positioning them as key hubs (Fig. 1H). Interestingly, while this network validated our multi-layered heuristics in selecting a predicted integrated core of evolutionary targets, it also indicated, through the co-expressed genes specific to each candidate that map onto different biological domains (Fig. 1I), that perturbing this set of targets could pinpoint genetic interactions between different domains of neurodevelopment regulation.

### Individual perturbations uncover requirements of KNL1, SPAG5 and KIF15 and dosage sensitivity of ADSL and BBIP1 in corticogenesis

Next, we designed multiple guide RNAs to perturb each gene and made two separate libraries to independently validate guides for up- and down-regulation. We then infected our CRISPRa line with the vector encoding SaCas9 and the two respective libraries of sgRNAs alongside non targeting control guides (NTC), generating one pool of lines bearing one saNTC and our compendium of spPerturbations, and a second pool expressing one spNTC next to our set of saPerturbations, see Fig.2A, Fig.S2). Mosaic organoids confirmed detection of all possible single-target perturbations (Table S1) and showed the expected morphology and markers distribution (Fig. 2B). To prioritize the two best guides per gene for each direction of perturbation we developed a ranking system that combines MixScape score (where a cell is considered perturbed if it shows significant transcriptional changes with respect to unperturbed cells, see Methods) with the effective detection rate of the perturbed gene, measured as percentage of cells expressing it, with *delta* calculated with respect to NTC cells marking the increase or decrease in the number of expressing cells ( Fig.S3 A)). From these scores, we generated 240 different combinations (Table S2) carrying out 2 separate lentiviral infections for 2 differentiation batches per infection (Fig. 2E), pooling 10 organoids per dish and profiling them through 7 separate sequencing runs (each containing a pool of cells coming from the 10 organoids). This approach maximizes averaging across conditions, ensuring a largely homogeneous distribution of perturbations across sequencing runs/pools, and hence data robustness, as shown by integration with Harmony (see Methods), with data clustered by cell cycle stage and sequencing runs homogeneously distributed across Leiden clusters ( Fig. S3 B-C).

**Figure 2.**
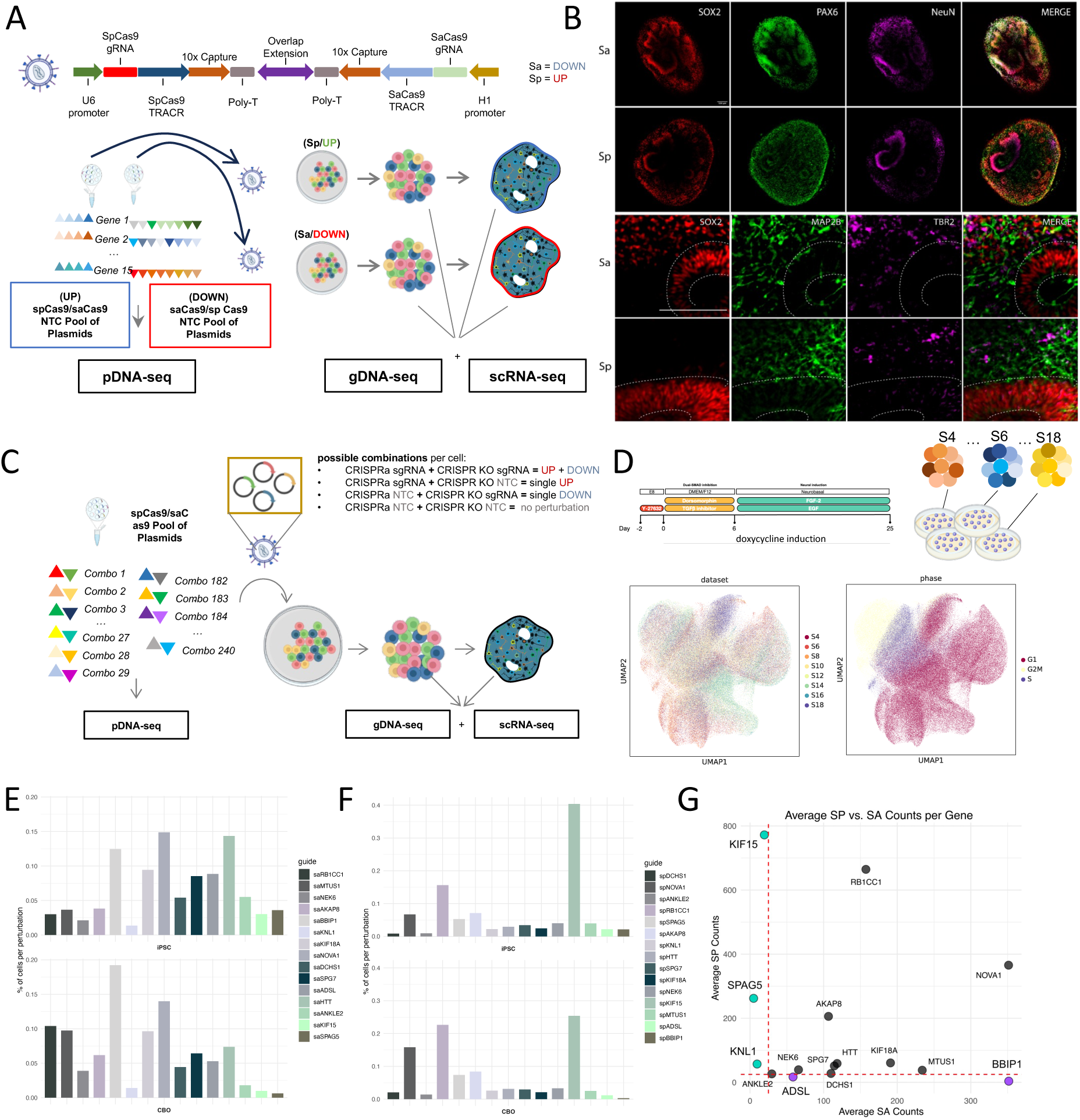
Guide selection and individual perturbation dropout screens. A) Experimental design for sgRNA guide selection and schematics of the construct, with the two bacterial Cas9 promoters (U6 for SpCas9 and H1 for SaCas9), trans-activating CRISPR RNAs (TRACR), 10x capture sequences for sgRNA detection, poly-T sequences and one overlap extension sequence to collate the Sa and Sp units. Two separate plasmid libraries were generated to insert sgRNAs for up- or down-regulation alongside a NTC, and distinct infections were carried out to induce single gene up- or down-regulation in iPSC, which were subsequently differentiated into mosaic CBOs. Four and eight guides per gene were tested for up-regulation and down-regulation, respectively. Plasmid, genomic and transcriptomic materials were sequenced to assess the cloning efficiency and targeting activity of sgRNA. B) Immunostainings of key early cortical markers reported for both up- and down-regulation mosaic CBOs, with SOX2 and PAX6 for ventricles and early radial glia progenitors, TBR2 (EOMES) for intermediate progenitors, MAP2B and NeuN for mature neurons. C) Experimental design of the final experiment. With 15 genes and two sgRNAs per construct, we estimated N*N-1 combination of guides (240 in total). D) Schematics of organoids differentiation protocol (above) and UMAP (below) obtained from 7 sequencing runs performed on a pool of 10 organoids per plate. Integration of sequencing run by Harmony shows mixing of runs and clustering by cell cycle on UMAP. E) Barplots depicting the percentage of cells per perturbation in iPSC (upper panel) and CBOs (lower panel). For each gene (x-axis), down-regulations are shown on the left and up-regulations on the right. G) Scatter plot of average high-quality cell counts per perturbation. The x-axis represents down-regulated (SA) counts and the y-axis up-regulated (SP) counts. Genes highlighted in blue (SA) and purple (SP) show significantly lower cell counts, indicating near-lethal perturbations.

Comparing (sa)guide abundances in iPSC and CBOs, we observed a differential representation during early corticogenesis for several perturbations (e.g., saBBIP1 increasing, and saHTT decreasing in CBOs). Examining *sp*-guide abundances, we observed a relative increase in representation of spNOVA1 and spRB1CC1 at later stages (Fig.2F-G). We thus calculated the number of cells bearing a specific perturbation, initially regardless of the detection of the concomitant paired perturbation, to maximize the signal derived by each of them (see Methods). Doing so allowed us to identify strongly depleted perturbations in CBOs (Fig. 2H, Fig. S3 D): the upregulation of *ADSL* and *BBIP1* and the downregulation of *KNL1*, *SPAG5* and *KIF15* are virtually incompatible with cortical organoidogenesis.

### Detecting gene interactions through bidirectional perturbations

Having validated our system for the yield of sufficient amounts of cells, perturbation, and cell type, we implemented a multi-step approach to define the regulatory relationships among our targets, from differential expression analysis to trajectory deconvolution, gene-regulatory network reconstruction and analysis (Fig. 3A).

**Figure 3.**
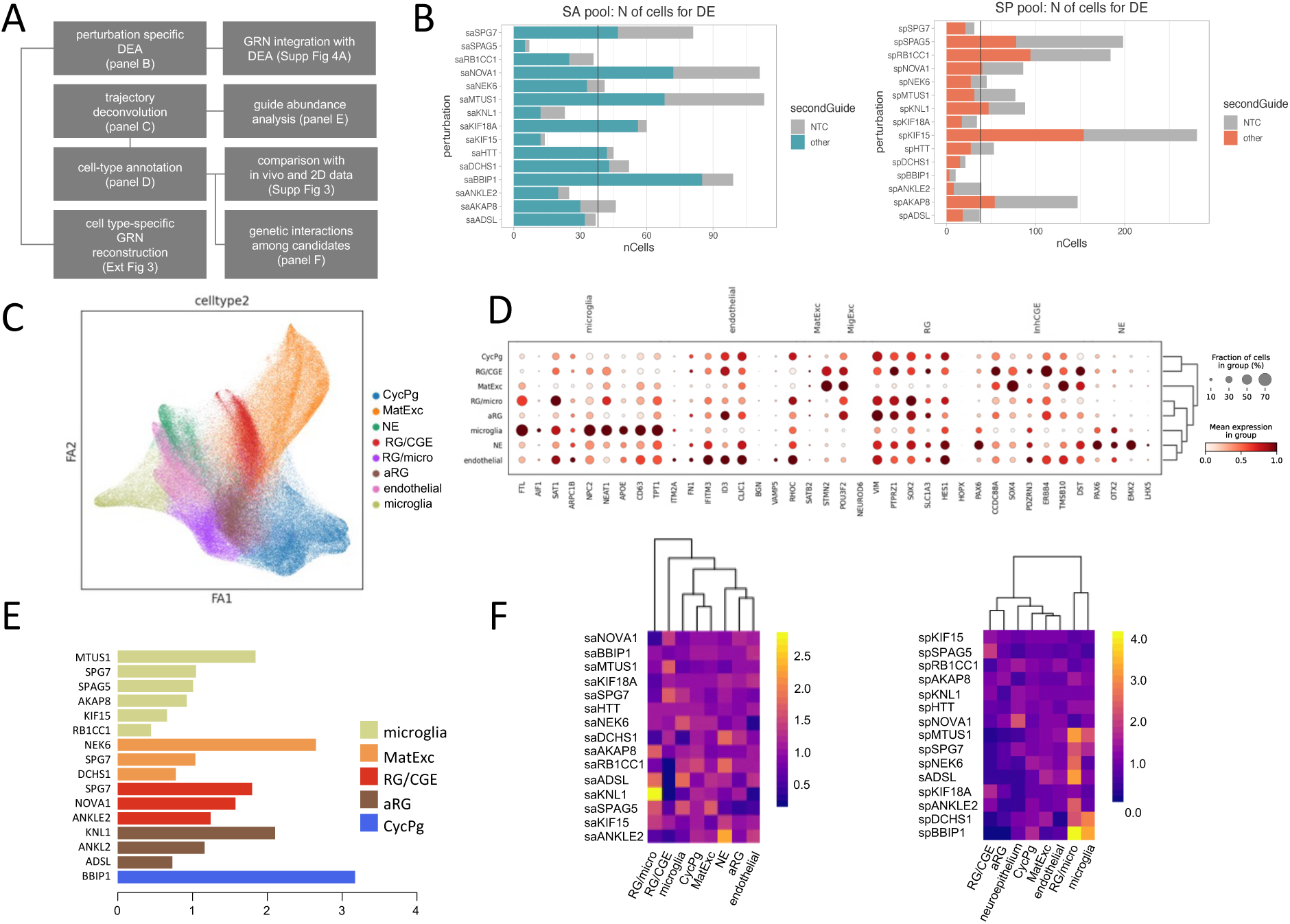
Deciphering cell type specific activity and gene interactions. A) Schematics of the analytical framework illustrating the sequential order and connections among analyses and the figures summarizing them; B) Stacked barplots of the number of cells in which a specific perturbation was detected. Grey bars represent cells with a single perturbation paired with an NTC, while colored bars report cells where the perturbation indicated on the y-axis is paired with a non-NTC. Light blue and light red bars correspond to the down-regulation and up-regulation pools, respectively; C) Drawgraph of the full dataset of 8 CBOs sequencing runs for a total of ∼250,000 cells. Each color indicates a different cell type: cycling progenitors (CycPg, blue), maturing excitatory neurons (MatExc, orange), neuroepithelial cells (NE, dark green), radial glia expressing caudal ganglionic eminence markers (RG/CGE, red), apical radial glia (aRG, brown), radial glia cells transcriptionally close to microglia (RG/micro), endothelial cells (light pink), microglia-like cells (light green); D) Dotplot of cell type-specific markers identified by intersecting top fetal cortex markers with the most expressed genes in each Leiden cluster; E) Barplot of cell type-specific enrichment scores (x-axis) measured by comparing the percentage of cell type markers found as target of each of our core genes with the percentage of markers of that cell type expressed in cells of that cell type. Significantly enriched core genes are displayed and color-coded by cell type (y-axis). F) Heatmap of cell type-specific guide enrichments measured as the percentage of cells bearing one specific perturbation (y-axis) with respect to the percentage of cells bearing the same guide on the full dataset. Cell types are indicated on the x-axis and perturbations (y-axis) are ordered depending on the number of core genes found to be differentially expressed in each of the single perturbations.

Previous work indicated ∼30 cells per perturbation as the minimal number to draw robust conclusions from scRNAseq CROP-seq experiments (31). Given the high complexity of our system, to gather robust target calls for each of our candidates, we started by pooling all cells with a given perturbation (e.g. upregulation of *ADSL*), independently of the companion perturbation, and performed differential expression analysis for perturbations represented by at least 40 cells, across cell types and sequencing runs (Fig.3B, see Methods). This equipped us with enough cells to describe the majority of perturbations with high confidence uncovering the targets of each gene while integrating cell type specificity and averaging the effect of companion perturbations and potential batch effects.

Specifically, we performed canonical dimensionality reduction and clustering, followed by annotation against fetal scRNA-seq data (*21*) (Fig. S4 A-C, see Methods). In contrast with the unperturbed controls, our double-perturbation mosaic organoids show a more variegated composition (Fig.3C), pointing to the impact, on several lineages of the interactions among our targets. In addition to the same (expected) cell types of unperturbed organoids, we observed less expected (for cortical organoids at such stages) the presence of early *EMX2*+/*OTX2*+ neuroepithelial cells, AIF1+ cells expressing microglia markers, *ITM2A*+ endothelial cells and two extra populations of cells expressing radial glia markers: one also expressing interneuron precursors markers, and one bearing an intermediate transcriptional makeup between CycPg and microglia-like cells, which we labeled RG/CGE and RG/micro, respectively (Fig.3D, Fig. S5 A). Notably, while the presence of canonical interneurons is being increasingly recognized in dorsal telencephalon organoids (*10,22,32,33*) it manifests typically at much later stages, pointing to a lineage acceleration triggered by some of our perturbations.

Among these cell populations specific to the bidirectional perturbation setting, the only cell type that was never previously described in patterned cortical organoids is microglia. We thus further characterized the identity of this cluster by probing the transcriptome iPSC-derived microglia (34) and observing the overexpression of *BBIP1* and *MTUS1*(Fig. S5 B-C).

Next, to assess the importance of our candidates in defining each of the cell type we identified, we implemented a scoring function that weights the number of markers of each cell type affected by the dosage of each of our candidates (*cell type specificity score*). The most significant results included strong enrichments for MTUS1 in microglia-like cells, *NEK6* in MatExc population, *SPG7* and *NOVA1* in RG/CGE, *KNL1* for aRG, and *BBIP1* in CycPg (Fig. 3E, see Methods).

To decipher our candidate genes’ contributions to the acquisition of each cell type identity, while contextually evaluating their reciprocal dependencies, we counted how many candidates were directly regulated by each candidate, and ranked them accordingly. Then, we integrated this information with the enrichment of guides targeting each candidate in each cell type (*cell type specific guide enrichment* in Methods) (Fig. 3F). The combination of this ranking with guide enrichments provides an indication of which cell type is populated by each perturbation, and which candidates are controlled by each other, thus providing a complementary and sufficiently independent measure with respect to the scoring function described above, where each candidate was tested for its ability to affect the expression of cell type markers. Indeed, integrating the gRNA modality with the scRNA modality, we found a crucial coherence between the perturbations detected in a given cell type and the effect of each perturbation on that cell type. Considering that cell-guide associations are purely based on guide detection across cell types and batches, and that candidate-targets associations are based on differential expression analysis across pseudobulks of multiple cell types, this constitutes a key validation of the ranking of each candidate with respect to the others. For instance, as shown in Fig.3E-F, perturbations of a given candidate (e.g. spMTUS1) that regulates the markers of a given cell type (e.g. microglia) cluster coherently by cell type, and indicate that the top-ranking gene (*MTUS1*) not only controls the highest amount of markers of that cell type, but also affects the other candidates perturbed in that cell type. Moreover, this measure is independent from the number of cells detected with a certain perturbation, and ranks *AKAP8*, *KIF15*, *KNL1*, *RB1CC1* and *SPAG5* above the other candidates.

It is worth noting that once perturbations are ranked in such a way, we generally observe an opposite distribution between the down- and the up-regulated sets of enrichments, suggesting a dosage-dependent behavior of our candidates. Moreover, the direction (up/down) of the perturbation provides a more specific indication of the importance of a gene for a specific cell type. For instance, the observations that *MTUS1* upregulation enriches microglia-like cells and that *MTUS1* downregulation is depleted in microglia, jointly point to *MTUS1* expression dosage as key regulator of microglia.

### Cell type specific regulatory gene interactions

To derive from differential expressions performed in pseudobulks from each individual perturbation the regulatory logic underlying cell type identities, we resorted to CellOracle (see Methods). First, to build a base gene regulatory network (GRN), we assembled a large reference set of regulatory regions drawn from the integration of multiomics data from cell lines, fetal cortex and brain organoid data (see Methods) and manually added the putative targets of each perturbed gene (see Methods). Then, we reconstructed cell type specific GRNs and leveraged cell type specific co-expression to zoom in onto the regulatory relationships of our candidate genes with their upstream and downstream effectors. As proof of concept, we leveraged the reconstructed GRNs of microglia-like cells to chart the genetic interactions between the three candidates showing highest cell type specificity scores and guide enrichments: *MTUS1*, *BBIP1* and *DCHS1* (Fig. S6 A). This microglial subnetwork highlights shared as well as unique interactors of the three factors, which include *NHLH1* downstream of MTUS1, and *NFIB* under the control of DCHS1, in line with their role in different cell types, and bring out *EPHA3*, *BCAT1* and *DLGAP1* as shared targets, coherently with MTUS1 and DCHS1 being expressed in either iPSC-derived microglia or neurons, respectively (Fig. S5 B-C). Notably, BBIP1 targets included key cell cycle and proliferation genes such as *CDK1*, *DLL3*, *MKI67* and *PRC1,* coherently with its enrichment in CycPg and the virtual lethality of its overexpression.

For RG/CGE, the other population besides microglia-like cells that is rarely if ever identified at this stage in cortical brain organoids, we uncovered different genetic interactions. Here, the degree of shared targets between the 3 master regulators KIF15, KIF18A and SPAG5 is paramount ( Fig. S6 B), with KIF15 and SPAG5 sharing key markers of early neocortical differentiation, including FOXP2, ID1, SOX4 and SOX11, and also, crucially, another of our candidate genes: NOVA1.

### Bidirectional perturbations and genetic interactions

To systematically quantify the degree of interactions across all our candidate genes and identify cell type specific biologically meaningful sets of interactions, we looked collectively at shared targets across cell types. Several of our candidates exhibit cell type specific gene interactions, suggesting modularity and further evidencing their cell type specific functions (Fig. 4A). KNL1, for instance, shares targets only with DCHS1, and specifically in CycPg, while in a microglia-specific manner, BBIP1 shares targets only with DCHS1, confirming the opposing roles of these two genes in determining microglial and neuronal fates, respectively.

**Figure 4.**
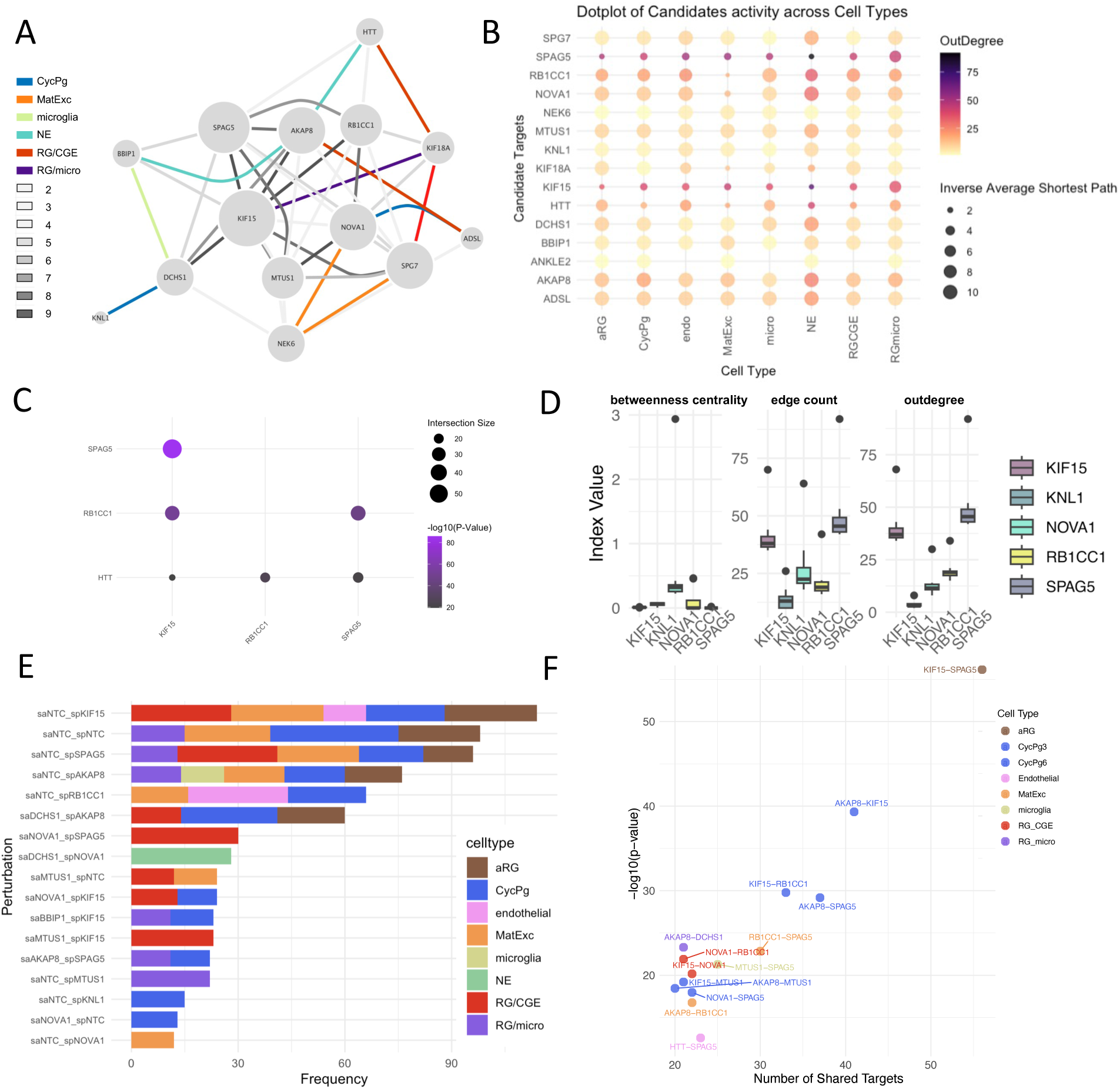
Cell type specific gene interactions among core genes and double perturbations. A) Graph representation of core genes interactions. An edge is drawn for all pairs of core genes sharing targets. Cell type-specific interactions are colored by cell type, with interactions shared across cell types indicated in grey and different grey shades indicating the number of cell types exhibiting the interaction; B) Dotplot of outdegree (number of regulated genes) and average shortest path (average distance between any two nodes in the graph, inverted to show higher values for smaller distances) for each core gene (y-axis) in each cell type-specific GRN (x-axis). Outdegree is reported by dot color, and inverse average shortest path values are reported as dot size; C) Dotplot reporting the number of shared targets across the four hubs identified in panel B, as KIF15, HTT, RB1CC1 and SPAG5. Dot size indicates the significance of the overlap measured by hypergeometric test, dot size reports the number of shared targets between core genes in each pair. X-axis and Y-axis report sets of gene targets, missing dots refer to redundant values and were removed; D) Box plot of betweenness centrality, edge count and outdegree scores reported for KIF15, KNL1, NOVA1, RBC1CC1 and SPAG5 (x-axis) across cell types (y-axis). Despite the high average shortest path, NOVA1 and KNL1 show high betweenness centrality and were thus included among the top core genes identified upon network analysis of cell type-specific GRNs; E) Stacked barplot reporting number of cells (x-axis) with exactly two guides detected (y-axis), colored by cell type; F) Scatter plot reporting the number of cell type-specific shared targets (x-axis) for each pair of core genes. Significance of the overlap (y-axis) was measured via hypergeometric test, using union of targets as background. Dots are colored by cell type.

Analyzing each cell type-specific GRN to rank candidates in terms of number of regulated targets (outdegree) we identified a small set that regulates several genes across cell types: *SPAG5*, *KIF15*, *HTT*. The combination of outdegree and average shortest path (a measure of the closeness of a node to any other in the network) allowed us to add RB1CC1 to this list and proceed with these 4 candidates as the most central ones (Fig. 4B). Examining shared targets and the significance of their intersections we found that *SPAG5*, *KIF15* and *RB1CC1* share a large significant number of targets, while *HTT* controls a different group of genes (Fig. 4C). Finally, to further gauge the connectivity of our core genes and identify potential hubs we measured betweenness centrality across cell types (in each cell type-specific GRNs) and found *NOVA1*, *RB1CC1*, and *KNL1* to be the top mediators (Fig. 4D).

Looking next exclusively at cells bearing two guides and specifically comparing the abundance of double candidate combinations vs single candidate-NTC combinations, and NTC-NTC combinations, we found all up-regulations and a vast majority of down-regulation, highlighting that almost all guides and individual perturbations are well represented while specific combinations of candidates decrease fitness (Fig. S7 A). Notably, the two missing downregulations (*HTT* and *KIF15*) are for genes whose upregulation is particularly abundant and equally represented across cell types, which points to a key minimum dosage dependence of proper cortical development for *HTT* and *KIF15*.

These observations acquire further significance when we analyze the distribution of double perturbations in a cell type-specific fashion. Considering only cells with two sgRNA guides detected, unperturbed cells are found exclusively in CycPg, radial glia populations, and MatExc neurons, confirming our earlier observation that our engineered line devoid of perturbation guides generates typically developing CBO. The upregulation of KIF15 shows a slightly higher complexity, expanding the cell types to RG/CGE populations and endothelial cells, which are usually found at later stages of CBO differentiation. Notably, specific double perturbations were found only in specific cell types such as, for instance: i) RG/CGE for either the downregulation of *NOVA1* coupled with the upregulation of *SPAG5*, or the downregulation of *MTUS1* coupled with the upregulation of *KIF15* (Fig. 4E); ii) neuroepithelial cell (NE) enriched by *DCHS1* downregulation coupled with *NOVA1* upregulation. Notably, in this analysis, the upregulation of *MTUS1* is found enriched in the subgroup of progenitors transcriptionally closest to microglia-like cells, which may thus be capturing the closest multipotent state preceding commitment to microglia-like state.

Looking at the genetic interactions captured by our GRNs, we found shared targets of NOVA1 and SPAG5, like *NFIA*, which is under the control of *EGR1* and *MEIS2* in RG/CGE, both of which are affected by - and directly correlated with - *NOVA1* dosage (Fig. S8 A). In the same cell type, we observe also a direct link between *MTUS1* dosage and *MEIS2* expression, which affects *NFIA*, and we observe an opposing tension between the activating effect of *MTUS1* dosage and inhibitory effect of *KIF15*, despite the fact that they both contribute to the inhibition of *NES* (Fig. S8 B). Collectively, these two subsets of the RG/CGE GRN reveal an indirect interaction between the two couples (*SPAG5*/*NOVA1* and *MTUS1*/*KIF15*) through their effects on *NFIA* and *MEIS2*, with a largely contrasting activity by MTUS1 and KIF15 that is unmasked thanks to the decoupling enabled by our double perturbation approach. In NE cells instead, we observe two largely independent activities of DCHS1 and NOVA1, with a few key shared targets. Here, we see again that one of the two genes (here *NOVA1*) acts on the targets of the other (*DCHS1*) through *NFIA* and *MEIS2*, while we unmask the synergistic activity of the two perturbed genes on *FAM181B*, *FGFBP3* and *SFRP2* (Fig. S8 C).

Building on these key cell-type specific interactions, we could then systematically interpret the interactions between our candidates (Fig. S9 -10). For this we quantified the number of shared targets between candidates, observing a strong genetic interaction between *KIF15* and *SPAG5*, which was not immediately obvious in the previous analysis on RG/CGE. We observed that these two factors act on the same set of genes already at the aRG stage (Fig.4F). Given its interaction with several candidates, along with the fact that saNOVA1 is the downregulation condition that is most abundant when coupled with upregulations of our candidates we characterized by GO enrichments the NOVA1 cell type-specific targets found in double perturbations. Only three interactions generated significant enrichments. KIF15 and SPAG5 showed coherent top categories related to translation, while AKAP8 showed enrichments for DNA metabolism, DNA duplex unwinding and cell division (Fig. S7C).

### Evolutionary outlook: linking potential candidates with the perturbed ones

In order to contextualize the nature of our perturbed candidates from an evolutionary perspective, we examined if their targets returned statistically significant overlaps with lists made of genes associated with signatures of positive selection (*24*) (Fig. 5A), genes that have accumulated an excess of high frequency mutations relative to their length and relative to other hominins (*9*) (Fig. 5B), and genes harboring virtually fixed high-frequency missense mutations in sapiens not found in other primates (*27*) (Fig. 5C). Taking advantage of our double perturbation logic, we could distinguish constituents of these lists based on their sensitivity to an increase or decrease in dosage of each perturbed gene. These overlaps suggest that *HTT* and *SPAG5* upregulation systematically affect genes under positive selection. Moreover, we found several of our candidates affecting genes that accumulated an excess of mutations with a clear direction specificity. Each of our candidates affected these excess mutation genes upon down- or up-regulation, highlighting the importance of gene dosage and paving the way to future works aimed at understanding whether derived mutations in modern humans increase or decrease the dosage or activity of the mutated genes. Finally, among our candidates, only *RB1CC1* significantly enriched genes that harbor fixed mutations in *Homo sapiens* with respect to other primates, suggesting a human-specific activity for this gene. None of these lists were significantly overlapping with targets of double-perturbations. Thus, we moved to genes bearing non coding HFV residing in their cortex-specific regulatory regions (see Methods), to understand the potential impact of our candidate genes interaction on development. Only six coupled perturbations significantly overlap with this list of regulatory relevant mutations (Fig. 5D).

**Figure 5.**
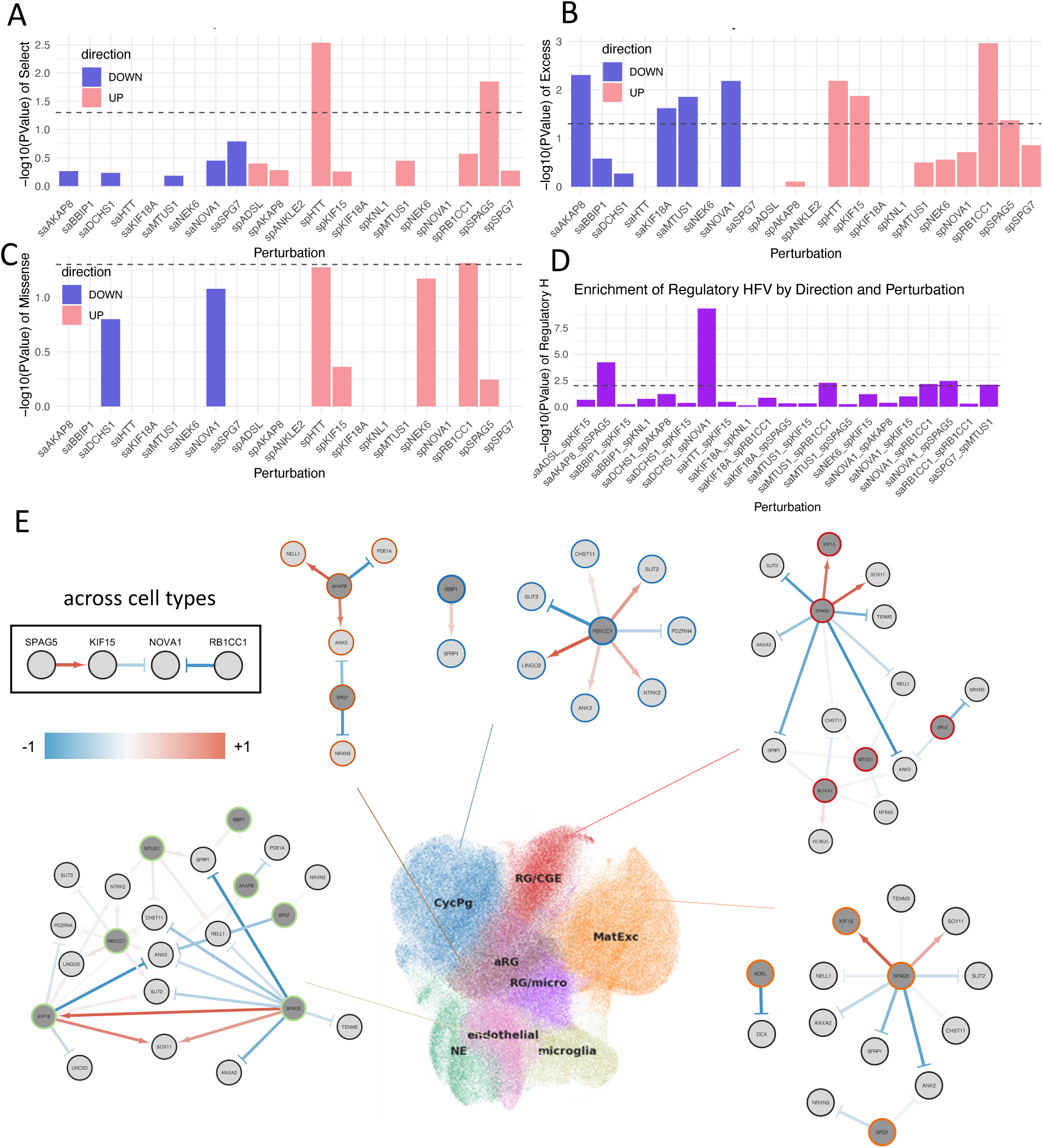
Ramifications of our core genes to evolutionary relevant gene sets. A) Barplot of significance values measured by hypergeometric test, comparing lists of genes affected by each perturbation and the list of genes associated with signatures of positive selection; B) Barplot of significance values measured by hypergeometric test, comparing lists of genes affected by each perturbation and genes that accumulated an excess of high frequency mutations relative to their length in *Homo sapiens* with respect to other hominins; C) Barplot of significance values measured by hypergeometric test, comparing lists of genes affected by each perturbation and the list of genes harboring virtually fixed high-frequency missense mutations found in *sapiens* but not in other primates; D) Barplot of significance values measured by hypergeometric test, comparing lists of genes bearing noncoding HFV residing at their cortex-specific regulatory regions; E) Graphical representation of networks for gene interactions found across all cell types (reported within the squared black box), cell type-specific GRNs involving our core genes and genes found in the lists used for panels A to D. Each GRN is linked to its specific cell type with a colored stick on the UMAP colored according to cell type. Cell type names are reported on top of each cluster.

Following up on the underlying logic of these overlaps, we extracted cell type specific GRNs involving our candidates and genes drawn from the human evolution-relevant lists we overlapped to explore the ramifications of our observations, and we isolated a series of small cell type specific networks of our candidates (Fig. 5E). First, we observed that certain candidate seems to affect different set of genes, in different cell types, allowing us to extend a few of them cross-regulate themselves across cell types, with *KIF15* as a direct positive target of SPAG5, and both KIF15 and RB1CC1 negatively regulating *NOVA1*.

## Discussion

Here we spearheaded dual CRISPR-Cas9 perturbations at single-cell resolution with guide capture multiomics to alter the expression and decouple the activity of a set of genes implicated in the evolution of the modern human cortex. Our approach builds on the growing recognition that complex traits, such as divergent brain ontogenies, require multiplexed genetic perturbations to make headway into the complex functional interactions among coding and non-coding elements (*35*). “Set-of-genes” approaches, whereby a selection of genes implicated in a given phenotype are perturbed in parallel to chart their individual contributions, have recently been applied to a range of neurodevelopmental conditions (*36–41*). By sampling the full space of mutual perturbations in all possible combinations, our experimental setup demonstrates the feasibility of experimentally dissecting the regulatory logic within such gene sets, and of doing so for multiple cell lineages in complex developing systems. Furthermore, in bypassing the effect of specific single-nucleotide variants and focusing on directional changes of evolutionary targets, our approach complements the work on paleoepigenetics and differentially methylated regions (*42*), with the crucial added specificity afforded by brain organoid models.

Our key insights can be summarized as follows. First, we found that two genes, KIF15 and SPAG5, whose modulation steers the proliferative activity of early progenitors, are at the top of the hierarchy among our targets, in terms of effect of their downregulation, average connections across cell types and guide enrichment of their upregulation across cell types. Both genes also proved capable of controlling a significant and large portion of genes bearing an excess of mutations in *Homo sapiens*, and specifically SPAG5 interactors showed a significant enrichment for genes under positive selection. Among the other targets, ADSL stands out for its inhibition of *DCX* expression in maturing excitatory neurons, pointing to a role of metabolic relay in setting the pace of neuronal maturation, in line with the role of *ADSL* in regulating glycolysis (*43*) and its promotion of autophagy (44), for which we recently uncovered a key role in the evolution of neuronal tempo (*16*).

Second, by reconstructing regulatory hierarchies beyond canonical TF-gene associations, we aimed at integrating the cytoplasmic and the nuclear partitions along which the regulation of cortical development has traditionally been tackled. Our results predict *KIF15*, *NOVA1*, *RB1CC1* and *SPAG5* to have had the most impact. In particular, the co-expression of *RB1CC1* with lysosomal genes, and its dosage impact on neuronal and RG/CGE markers, points to a key role in neuronal differentiation, converging with the recent discovery of CHD2 activity in pacing development, differentiation and activity of cortical neurons by modulating the lysosomal pathway (*16*). Moreover, while the roles of *DCHS1* and *NOVA1* in evolution have so far been studied in isolation (*11,45*) our work highlights their synergy, partitioning it into two mechanisms, one featuring a small set of key downstream targets and the other entailing a lateral flow circuit whereby NOVA1 acts on DCHS1 targets through the NFIA and MEIS2 mediators. An ancestral isoform of DCHS1 featuring increased affinity for EPHA4 has been shown to steer the evolutionary rearrangement of neocortex-striatum ratio (Cappello and coworkers), converging with our finding of a direct effect of DCHS1 on EPHA3 and highlighting the complementarity of the two model systems in capturing coordinated regulation of ephrin receptors by DCHS1 as an exquisitely dosage-sensitive mechanism in the evolution of the modern cortex also vis a vis subcortical fates.

Third, the juxtaposition of multiple perturbations allowed us to decouple the activity of SPAG5, KNL1 and KIF18A, which were until now predicted to have played a largely shared role in evolution (*5*). Fourth, the dominance of glial fate and ciliogenesis markers among several of our perturbations points to the relevance of signals from the ventricles which also affect astrocytes differentiation, in line with independent work associating neurocranial globularity with ventricle and white matter pathways (*46*).

Finally, our set-up also led to the identification of genes steering early cortical development that might be repurposed to develop new protocols for the streamlined generation of more complex cortical organoids. For instance, the finding that *MTUS1,* which is expressed in iPSC-derived microglia, actually determines the acquisition of such fate upon overexpression in the context of a patterned cortical organoid, provides new inroads for the generation of this cell type in vitro, eventually by seeding mosaic organoids bearing optimal amounts of cells harboring this perturbation.

## References and Notes

### Limitations of our study

While our study paves the way to the combinatorial dissection of evolutionary targets at scale, also in combination with our recent advances in brain organoids multiplexing (22) the number of our core genes remains necessarily modest with respect to large scale CRISPR-Cas9 screenings focused on single perturbations. In this respect, based on our implementation of dual perturbation systems with the a priori calculation and ex post assessment of the optimal number of genes to target given the complexity of brain organoids single cell readouts (fifteen), we estimate that with reduced costs of sequencing and further optimisation in library preparation and guide detection, it should be possible to perturb up to 25-30 genes in a comparable setting. The network size reconstructed on the basis of our perturbations includes 370,000 interactions, 220 regulators and 3190 target genes. This number depends on the capabilities of the GRN reconstruction method that we used, and, with greater computational power, we expect this could be doubled in the near future. We also inherit from paleogenomics the inherent limitations derived from the small number of high-coverage genomes of extinct hominins, leaving our target gene criteria subject to future modifications. However, we implement the use of high-frequency variants also in the attempt to compensate for this imbalance. Our study is inevitably confined to the effects of perturbations in cerebral cortex organoids. However, precisely the comparison of our results with those from unperturbed organoids, as well as of unperturbed cells within mosaic double perturbed organoids, allowed us to pinpoint the appearance of cell types that are not typically present in this organoid system, extending the reach of our conclusions and proving the value of this approach also for synthetic biology inroads towards organoid refinement. In the future, other brain regions, such as the cerebellum, will benefit from being examined with the same methodology. Since we did not introduce specific derived mutations in our experiment, in light of the direction specific effects of perturbations we uncovered here, future work might be focused on determining the impact of such mutations on gene dosage and protein activity, to extend our conclusions and validate them in terms of evolutionary variants’ specificity. While work complementing ours already allows us to infer the effect of several mutations (e.g., the mutation affecting ADSL) (14,47), much more work is required in this area, particularly for genes harboring several nearly fixed missense mutations, such as SPAG5 or KIF15. Likewise, here we focused on a comparatively early time point to combine organoids complexity with experimental tractability for cell type specific assessments. However, when focusing on the exact effect of the relevant mutations, they should be studied over multiple timepoints and a longer differentiation time should be the next frontier. There is clear evidence from genes implicated in neurodevelopmental disorders (48,49) that mutations have different functional effects at distinct developmental stages, and we expect this to be true for the evolutionary variants of at least several of the genes studied here, in light of their effects across cell populations on the downstream pathways we identified.

## Resource availability

Raw data will be available upon request.

## Acknowledgments

We thank present and former members of G.T.’s laboratory for insightful feedback provided on this study. We also thank the Flow Cytometry and Imaging facilities of Human Technopole and the European Institute of Oncology (IEO). We thank the European School of Molecular Medicine (SEMM) at which A.S., S.T., O.L are/were enrolled as students for their PhD degree program in Systems Medicine. We thank Silvia Cappello for fruitful discussion. Some components of the schematics were adapted from BioRender. The project, carried out in G.T.’s laboratory at the IEO and at Human Technopole, has been funded by Horizon 2020 Innovative Training Network EpiSyStem (GT), H2020 Marie Skłodowska-Curie Actions, Grant/ Award Number: 765966 (AS). C. B. was supported by Project PID2023-146627NB-I00 (Spanish Ministry of Science, Innovation, and Universities CIENCIA/AEI), project 2021-SGR-313 (AGAUR/Generalitat de Catalunya) and a Leonardo Grant for Researchers and Cultural Creators, BBVA Foundation. A.S was funded by the Leverhulme Trust under a Study abroad studentship. A.V. was supported by Telethon Research Grant GGP19226 (GT) and RE-MEND (101057604).

## Author contributions

Conceived the study : G.T., C.B. and A.V.;

Developed the project’s design and implementation G.T., C.B., E.V., A.V., A.S. and M.B.;

Conducted experiments A.S, O.L, M.F.P & S.T;

Analyzed data A.V, V.F, O.L, F.P., J.M., D.C., E.C.V. & C.B.;

Planned experiments A.S., A.V., S.T., E.V. C.B. & G.T.;

Designed the bidirectional perturbation strategy M.B., A.S., A.V., C.B. and G.T.;

Interpreted the data and wrote the manuscript with input from all authors A.S., A.V., O.L., C.B. & G.T.;

Supervised the study A.V., C.B. & G.T. Experiments were performed in G.T. laboratory.

## Supplementary materials

Materials and Methods

Figs. S1 to S10

Tables S1 to S2

References (50-64)

## Materials and Methods

### Gene candidate selection

Our primary source of comparative data (*9*) catalogs segregating sites between *Homo sapiens* and high coverage genomes from two Neanderthals and one Denisovan individuals (*1–3*), which we supplemented with data from an additional high-quality Neanderthal genome (*4*) published since. Following the criteria at the basis of our starting catalog (*9*), allele ancestrality was determined based on publicly available multiple genome alignments (*50*) or, in case this information was not available, from the macaque reference genome (*51*). Allele frequency was determined from the dbSNP database build 147 (*52*). A derived allele was deemed “highly frequent” if it occurred globally across modern human populations above a 90% threshold. Lists of fixed protein-coding changes employing a range of comparative considerations and filters (*5,11,27*) were also consulted.

### Cell reprogramming

KOLF2C1 line was purchased from the Wellcome Trust Sanger Institute and was reprogrammed into iPSCs using the Sendai virus. Line reference name is HPSI0114i-kolf_2-C1.

### hiPSC maintenance

hiPSC were cultured in TeSR-E8 medium (Stemcell technologies, 05990) supplemented with penicillin-streptomycin (P/S, 100 U/mL; Thermo Fisher Scientific, 15140-122), with daily media change, at 37 °C, 5 % CO2 and 3 % O2 in standard incubators. hiPSC were grown on matrigel-coated dishes prepared as follows: matrigel stock solution (Corning, 354248) was diluted 1:40 in DMEM /F-12 1:1 medium (Lonza, BE12-614F and Thermo Fisher Scientific, 11765054, respectively) supplemented with P/S 100 U/mL and used to coat dishes for 15-30 minutes at 37 °C. Passages 1:6-1:8 were performed using ReLeSRTM (Stemcell technologies, 05872) or Accutase solution (Sigma-Aldrich, A6964). ReLeSR was used to detach hiPSC in clumps for expansion and standard maintenance. Accutase solution was used for single cell passaging; in this case, ROCK inhibitor 5μM (Sigma, Y0503) was added to the culture overnight (ON) to enhance single hiPSC survival. Cryopreservation of hiPSC was performed by single cell dissociation and storage in complete TeSR-E8 medium plus 10% DMSO supplemented with ROCK inhibitor 5μM.

### Cerebral cortical brain organoid generation and maintenance

Cortical organoids were generated using an adaptation of the previously described protocol published by Pasca lab (*53*). hiPSCs were expanded on matrigel-coated 10 cm plates and dissociated at 60% confluency with Accutase solution for 3 minutes. Cells were centrifuged to remove the enzymatic suspension (160 Gs for 3 minutes). After resuspension in TeSR-E8 medium supplemented with 5 μM ROCK inhibitor cells were counted with a TC20 automatic cell counter (Biorad) and seeded into 96 ultra-low attachment well plates (S-bio Duotech, MS-9096UZ) at a final concentration of 2 x 104 cells in each well. EB are generated by single-cell aggregation; therefore, plates were centrifuged at 160g for 3 minutes to enhance EB formation. The following day medium was not changed, leaving EB undisturbed. At this point, Dorsomorphin (5 μM, MedChem express, HY-13418A) and TGF-β inhibitor SB-431542 (10 μM, MedChem express, HY-10431) were used to perform dual-SMAD inhibition, pushing neuroectoderm specification. Dual-SMAD inhibition was kept for a total of 5 days, with daily media change. On day 6 the differentiation medium 2 was added until day 25 with daily media change for the first 12 days, and then every other day. The differentiation medium 2 was composed of neurobasal medium (Thermo Fisher Scientific, 12348017) supplemented with 1X B-27 supplement without vitamin A (Thermo Fisher Scientific 12587001), 2 mM L-Glutamine, P/S, 100 U/mL, 20 ng/mL FGF2 and 20 ng/mL EGF (Thermo Fisher Scientific, PHG0313). Human FGF2 and EGF were used to amplify the pool of neural progenitors. On day 12, organoids were moved to ultra-low attachment 10 cm dishes and grown on shakers to enhance oxygen and nutrient supply.

### sgRNA selection process

CRISPick (*54,55*) helps rank and select the best CRISPRko/CRISPRa sgRNA sequences. It prioritizes sequences that work well while avoiding off-target effects

### Engineering of the CRISPRa system in hiPSCs

The CRISPRa system was engineered in the wild-type pluripotent stem cell line KOLF2C1 (which is registered in the European Human Pluripotent Stem Cell registry, is already characterised comprehensively including in terms of its whole genome sequence) to express SP-dCas9-VPR with doxycycline-inducible expression (PB-TRE-dCas9-VPR was a gift from George Church, Addgene plasmid #63800) (*56*), which is fused in tandem to the VP64, p65 and Rta (VPR) transcriptional activators for efficient gene activation upon sgRNAs transduction (*54*). The Piggybac vector was transfected into the KOLF2C1 cell line via electroporation (Neon Transfection System). The mix prepared before electroporation consisted of the CRISPRa PB Vector (2.5ug plasmid DNA), transposase (250ng), 500,000 cells and completed with Buffer T (NEON transfection Kit) to 120ul. Once the mix was prepared, it was electroporated under the following conditions: 900V, 20ms and 3 pulses. It was subsequently plated in a 100mm plastic plate, in TesR E8 + Ri (200x). Successfully transduced cells were selected with hygromycin (200 μg/mL) and single clones were expanded for 14 d. A total of 20 clones were picked, whilst the rest of the plate was polyclonally frozen. The 20 clones were expanded and frozen; only three clones were chosen to validate the efficiency of dCas9 incorporation. Clones were picked under the microscope and transferred to a 30mm plastic plate. Doxycycline was added at 2ug/ml, for 3 days, 5 days and 7 days respectively.

### Validation of the CRISPRa system in iPSCs and day 25 cortical organoids

To test functionality of the expanded clonal CRISPRa line, we first performed a western blot for Cas9 to evaluate the expression level of the protein in doxycycline induced cells. Once that was confirmed we then transduced cells with sgRNAs to activate the imatinib efflux transporter ABCG2 (5′-GCCACTGCGTTCAGCTCTGG-3′). The library vector sgLenti (sgLenti was a gift from Mike McManus, Addgene plasmid #105996) (*17*) was prepared by restriction digest with AarI (Thermo Fisher) at 37 °C overnight, followed by 1% agarose gel excision of the digested band and purification via NucleoSpin columns (Macherey-Nagel). Using a standard T4 ligation, phosphorylated ABCG2 sgRNA were annealed and ligated into the lentiviral backbone. The ligation was then transformed in STBL3 cells and plated the following day. 6 colonies were picked from the plate which were then mini-prepped and sent for sanger sequencing to validate that the cloning had been successful. Once it was validated, a 3rd generation lentivirus was produced which was introduced to the CRISPRa iPSC control line.

Clonal lines were screened for functionality of CRISPRa via flow cytometry analysis of >100,000 cells stained with CD338 (ABCG2) antibodies (Miltenyi, 130-104-960). unctional CRISPRa activity displayed stable function throughout 70 days of persistent doxy induction. All samples for mCherry positive-ABCG2-harboring sgRNAs were compared to the same vector harboring a Non-Target-Control sgRNA.

The same concept was introduced in cortical organoids (25 days after generation of cortical spheroids). Due to the 3D nature of cortical organoids, we decided to induce doxycycline from day 0 of differentiation of the CRISPRa line harboring the sgRNA for ABCG2, Doxycycline induction began on Day 0 of organoid generation, in 4ug/ml doxycycline, The overexpression was evaluated both via FACS analysis but also Western Blot.

### Library production

Pooled libraries for expression of single sgRNAs were made with oligonucleotide pools obtained from IDT (oPools) in conjunction with IDT Ultramers for the NTC sgRNAs (one per Cas9 organism). For cloning of the pools, oligonucleotide inserts were designed with 5′ BsmBI sites followed by 20 or 21 nt crRNA (depending on the Cas9 organism),82 nt tracrRNA, 10x genomics capture sequence and 17 nt complementary sequence. The oligonucleotides for SpCas9 sgRNAs and SaCas9 sgRNAs were separately mixed together at a concentration of 5 μM each. 10 μL of each pool of oligonucleotides was then combined in a 100 μL reaction and extended using NEBNext (New England Biolabs) with an annealing temperature of 48 °C. The resulting dsDNA was purified by spin-column then ligated into the BsmBI-digested pPapi vector (pPapi was a gift from John Doench and David Root, Addgene plasmid #96921)(*18*) using 80 cycles of Golden Gate assembly with 100 ng insert and 500 ng vector using NEB Golden Gate Enzyme Mix (BsmBI-v2) (*18*). The DNA was then transformed in STBL3 cells, which was then plated at a ratio of 1:1000, whilst the rest was added to LB broth and let to grow overnight. After <20 hours, the number of colonies found on the plates were indicators of the successful transformations that occurred within the broth. The LB broth was then purified via Maxi-prep, which yielded plasmid DNA containing the inserts of interest (saCas9 pool with 120 sgRNAs with the spCas9 NTC sgRNA found in the second position of the insert, and spCas9 pool with 60 sgRNAs with the saCas9 NTC sgRNA in the second position and finally a control with the two corresponding sgRNA NTC in the two positions). The vectors were then prepped for virus production. We note that individual constructs to express two sgRNAs can be constructed either by the overlap-extension of individual oligonucleotides (found in the oPools) or by the use of pre-synthesized double stranded oligonucleotides that can be directly cloned into the vector backbone.

### Plasmid DNA and genomic DNA preparation, sequencing and validation

Plasmid DNA (pDNA) was isolated using the mini-prep kit as per the manufacturer’s instructions. The concentration of these preparations was determined by UV spectroscopy (Nanodrop and Qbit). For the pPapi vector, dual sgRNA cassettes and plasmid DNA were PCR-amplified and barcoded with sequencing adaptors (Illumina TruSeq adaptors, TruSeq primer binding sites and indexes on the 5’ end of the oligonucleotide) using NEBNext. Amplified samples were then purified with 1% agarose gel excision of the digested band and purification via NucleoSpin columns (Macherey-Nagel). The purified band was then sequenced on a NextSeq sequencer (Illumina) with 50 nt paired-end reads. The same was done for genomic DNA (i.e. iPSC pools and cortical organoid pools); DNA was extracted via TRIZOL: From the plated transformation, 4 colonies were picked and purified via mini-prep; each colony should represent a specific sgRNA. The purified colonies were analysed via Sanger Sequencing. The purified pool of plasmids (i.e. the broth) was also analysed via sanger sequencing. The plasmid DNA of the three samples (saCas9 pool/spCas9 sgRNA NTC, spCas9 pool/saCas9 sgRNA NTC and spCas9 sgRNA NTC/saCas9 sgRNA NTC) were submitted to NGS to validate the successful cloning by understanding how many times each sgRNA existed within each pool.

To evaluate library representation and distribution of the sgRNAs in the pool, plasmid DNA was sequenced on an Illumina NovaSeq 6000 with 50 bp paired-end reads. A custom bash script was then used to count the occurrences of each sgRNA in the sequencing library. Briefly, reads either in R1 or R2 files (for Sp or Sa pools, respectively) were scanned for matches with the sgRNA sequences, without allowing mismatches. Due to our cloning scheme, only the first 16 nucleotides of the guides transcribed from the U6 promoter could be resolved, so the sequences of the sgRNA for SpCas9 were trimmed to generate 16nt barcodes. NTC were also counted in the same way. Unique hits were counted for each sgRNA to generate a table of raw counts, which was then imported in R for summary statistics and plotting.

### CRISPRa line infection and subsequent differentiation

iPSCs were infected with an MOI of 0.3, after the lentivirus was titrated via qPCR. After 48 hours of infection, the cells were selected with Puromycin at 2ug/ml, until the negative control cells had died. Since there are specific combinations within the cell pools, and no combinations were to be outnumbering any others, the cells were left in fresh medium for only 24 hours before differentiation into CBO began.

### Virus production

48 h before transfection, HEK293T cells were seeded in 150mm dishes at a density of 2×106 cells per dish in 15mL of DMEM + 10% FBS. Transfection was performed using a 3rd generation lentivirus production, as per usual. Briefly, the cells were transfected with the vectors suitable for 3rd generation lentivirus production, VSV-G (9ug), pMDL (12.5ug), pREV (6.25ug) and the sgRNA containing vector (pPapi, 32ug). The vectors were incubated at room temperature for 20– 30 min, during which time the HEK293T cells were replenished with fresh media. After this incubation, the transfection mixture was added dropwise to the surface of the HEK293T cells, where the plates were transferred to a 37 °C incubator overnight. The following day, the media was removed and stored at 4°C and replaced with fresh medium and left in the incubator for the rest of the day. After 24 hours, cells were harvested and ultracentrifuged at 40,000G at 4°C. The supernatant was discarded and the pellet was resuspended in 1% PBS.

### sgRNA library design for pre-screen and final screen

For the initial screens (CRISPRa pool with saCas9-sgRNA-NTC and saCas9 pool with spCas9-sgRNA-NTC), a sgRNA library consisting of 120 sgRNAs for one library and 60 sgRNAs for the other, respectively, was designed in order to find the best hitting sgRNA/gene. For the final screen, the process was mirrored, with the only difference being that we chose the two best hitting sgRNAs/gene and implemented them within the same backbone.

### Tissue clearing, immunofluorescence and data analysis

Cortical organoids were cleared and stained at day 25, using the MACS Clearing Kit (Miltenyi Biotec). First, organoids were fixed in 4% PFA without methanol for 20min and then permeabilized for 6h while shaking at RT. Samples were later immunostained with primary (for 48h) and secondary (24h) antibodies with five long washes in between to remove unbound antibody. Organoids were then embedded into 1% agarose (in ddH2O), and following solidification, the blocks were gently cut and transferred to 2 mL tubes. Dehydration using concentration gradients was performed for 24h and samples were later incubated in clearing solution for 6h. Samples can be put on hold at this point if stored in imaging solution. Samples were acquired using the Nikon spinning disk CSU-W1 with confocal configuration (Nikon, Japan).

### Single-cell 10x CRISPR-GEX library, quantification and quality control

Raw reads were aligned to Human genome assembly GRCh38, using *cellranger* v 7.1.0 from 10x Genomics. Gene Expression (GEX) modality was processed via Scanpy version 1.9.2. Good quality cells were selected by filtering out those expressing less than 300 features (genes) and features present in less than 25 cells. Normalization was performed following standard scanpy procedures, to correct for UMI count. Unperturbed organoids data was first analysed independently and not batch corrected. Perturbed organoids data was processed entirely, upon integration of sequencing run by Harmony (*57*). KNN was applied to identify cells neighborhood setting k = 50. A first step of Leiden clustering (resolution = 0.5) was applied to KNN neighbours identified on the first 10 principal components. MAGIC was applied to impute expression of canonical cell type marker genes and annotate cell types on leiden clusters. Clusters of cells whose marker genes were identified as ribosomal or mitochondrial via scanpy were removed, and normalization and clustering was perfomed again starting from the remaining cells’ raw data. UMAPs were calculated through scanpy, as well as diffusion maps. To generate a *drawgraph,* PAGA and ForceAtlas2 algorithms were applied to reconstruct differentiation trajectories as in (*24*). To annotate cell type we ingested the cortex specific cells from (*21*) on the single-cell RNA-seq data from (*58*) to obtain a large number of cortex cells from the former paper while bearing the more precise cell type annotation produced in the latter. This allowed us to extract the top 1,000 expressed genes of each cell type, and measure the expression of these genes in the leiden clusters of our perturbed organoid data. The intersection of the top 1,000 markers of fetal data with the top 20 genes expressed in each of our clusters, allowed us to attribute a cell type to each of them. Only a few clusters (later annotated as neuroepithelium and microglia-like cells) did not show clear markers of the expected cell types (neural precursors, radial glia and excitatory neurons), and were thus annotated based on their top expressed genes, including *EMX2* and *OTX2* for neuroepithelial cells, *AIF1* (IBA1 protein coding gene) cells for microglia, *ITM2A* for endothelial cells.

### sgRNA-cell association *in silico*

Given the peculiarities of our construct and the limited efficacy of the 10x paradigm, our work did not reach the promised levels of guide detection. This was confirmed both in the initial guide selection experiment and in the final experiment, both in iPSC and CBOs. Due to low capture efficiency, fewer cells containing the relevant perturbations passed the detection filter. This situation forced us to massively increase the number of cells/organoids to be sequenced (from the initial 30k cells target we set to profile 210k cells). Moreover, we resorted to identifying cell barcodes and sgRNA sequences in the raw fastq files before launching cellranger v 7.1.0, and identifying guides using the *grep* function of seqKit software, by accepting maximum 2 mismatch in the sgRNA sequence and none in the cell barcode. This avoided confusion between guides and certainty of cell barcode calling. We verified that no guides were found in the unperturbed organoids profiled with the 10x capture seq, and that sgRNA found by *cellranger* would exactly match the ones we associated to cells.

### Co-expression analysis

Following leiden clustering (resolution = 0.5, dividing the data into 10 clusters), unperturbed organoids single-cell were aggregated via pseudobulk by summing raw reads of cells of the same leiden cluster. This provided us with 10 pseudboulk RNA-seq samples. A regression analysis was thus performed using edgeR as in (*59*), to identify genes whose expression was correlated with each of our 15 core genes. The resulting logFC and -log10(FDR) of each analysis were multiplied, to obtain a score that was used to rank the top 100 genes co-expressed with each of our 15 core candidates. GO enrichment was then performed by topGO to estimate the function of our candidates. The significantly co-expressed genes (FDR < 0.05 and |FC| >= 1) were collated into a candidate-target table that was then imported into Cytoscape to generate the networks reported in Fig. 1H and Supp. Fig. 1E. For each of our 15 core genes, co-expressed genes and their candidate specific edges were coloured in *shocking pink*.

### sgRNA guide efficacy scoring

In the absence of tools for CRISPR guide assessment in Scanpy, the data was first converted and loaded on R with Seurat version 4. Each cell showing at least 2 reads per perturbing guide and associated with a unique guide was used for the purpose of guide ranking and expression detection of target genes. Mixscape (*60*) has been used to gauge guide efficacy in the single perturbation screening and calculate a Mixscape score for each guide. Given the well-known sparsity of single-cell RNA data, and the eventual inability of measuring fold-changes of perturbed genes, on top of Mixscape scoring we applied a further filter for guide ranking. Briefly, starting from the simple assumption that if a gene is more expressed it will be easier to detect it, we measured the percentage of cells in which the perturbed gene was detected to determine, by contrast with NTC bearing cells, if a gene had putatively been up- or down-regulated.

### Dropout screen of CRISPR perturbations

To identify lethal or nearly lethal perturbations, we counted the number of cells bearing each perturbation, summing cells whose perturbed gene and direction was equal albeit obtained through different sgRNA sequences. Then, we performed a multivariate variance analysis to verify whether a certain gene perturbation would significantly sit far from the distribution of all other perturbations. This indicated only 5 perturbations as significantly below the detection distribution: saADSL, saBBIP1, spKIF15, spKNL1 and spSPAG5, which all were found in less than 20 cells.

### Differential Expression Analysis on Perturbations

To identify the putative targets and genes affected by our 15 core genes perturbations, we aggregated all cells bearing a specific perturbation coupled with any other guide. Instead of pseudobulks, since our coverage target was high (50,000 reads per cell) and the number of cells with a specific perturbation could be highly variable, we resorted to differential expression through scanpy, performing *t-test,* through the *scanpy.tl.rank_genes_groups* function, for all perturbations represented by at least 40 cells. Significantly differentially expressed genes were considered “direct targets” of each perturbed gene in the following gene regulatory network reconstruction analysis.

### Differential expression Analysis on iPSC-derived microglia

We performed differential expression on data from GSE163984 with edgeR, using standard procedure, using *estimateDisp* with *robust =TRUE,* and *glmFit* function (*40,60*). We performed a single differential expression setting three groups: iPSC, iMG (iPSC-derived microglia) and iN (iPSC-derived neurons), to identify patterns of expression that would distinguish the three cell types. Only genes with FDR < 0.05 and |FC| > 2 were considered to filter our core genes’ cell type specific expression, and SPAG5 or MTUS1 targets (Fig. S4B-C).

### Gene regulatory network reconstruction

To build gene-regulatory networks and leverage the information provided by the perturbations we leveraged CellOracle v.0.10.15 (*61*). To build TF-gene annotations (*base GRN* in CellOracle) we first generated a repository of regulatory regions, collating data derived from single-cell organoids multiomics (*23*), Psychencode cortex specific enhancers (*62,63*), co-accessible peaks in fetal cortex development (*64*). These regions accounted for ∼450K region-gene associations, and 241065 genomic regions. This set of regions was provided to CellOracle to perform motif annotation and associate TFs to gene pairs. CellOracle allows to impute external information on gene regulation, in the form of a TF-target table, where each so called “TF” was one of our perturbed genes and the “target” genes were the genes differentially expressed upon the “TF” perturbation. Thus, we provided the outputs of our differential expression analysis on perturbations, to leverage CellOracle ability to measure cell type-specific relations between TFs and their targets based on co-expression. Cytoscape v3.10.2 was used to plot network subsets coming from GRNs.

### Master regulatory analyses

To refine our cell type annotation, we leveraged the GRN generated by CellOracle. First, we performed a cell type specific master regulatory analysis on these GRNs, by calculating the percentage of genes targeted by a certain TF in a specific cell type with respect to the percentage of genes targeted across cell types. Then, we calculated via hypergeometric test the overlap between the targets of each couple of TFs in each cell type, with respect to the union of their targets in each cell type. This provided us with cell type specific genetic interactions between TFs couples. For this analysis (Fig. S4A) we excluded our core perturbed genes.

To calculate the cell type specificity score of our core genes, we calculated an enrichment by dividing the proportion (%) of markers of a given cell type (identified previously in reference fetal data) targeted by each of our 15 core genes, by the proportion of markers of the same cell type expressed in the cells annotated accordingly. Significance of the intersection between targets of each candidate and markers of a given cell type was measured by hypergeometric test, and only significant enrichments were considered. As a complementary measure, we calculated the “cell type specific guide enrichment” which instead of focusing on the putative ability of a core gene to regulate the markers of a given cell type, focused on the frequency of each perturbation in each cell population. Hence, the cell type specific guide enrichment was obtained for each perturbation as a measure of the percentage of cells bearing a certain perturbation in a cell type divided by the percentage of cells bearing the same perturbation across all cells (independently of cell type identity).

### Minimal Network deconvolution

Starting from differential expression performed in each dataset we have identified the transcriptional effects directly affecting perturbed genes upon perturbation of one of the other target genes. Briefly, if a gene was upregulated upon KO of another gene, we considered the second gene as a putative inhibitor of the first. If a gene was downregulated upon KO of a certain gene, the second would be considered as an activator of the first. Similarly, mirroring rules were applied to parse the effect on our target genes upon up-regulation of one of the others. Python igraph library was used to represent all identified connections.

## Reference List

1. Meyer, M. et al. A High-Coverage Genome Sequence from an Archaic Denisovan Individual. Science 338, 222–226 (2012).

2. Prüfer, K. et al. The complete genome sequence of a Neanderthal from the Altai Mountains. Nature 505, 43–49 (2014).

3. Prüfer, K. et al. A high-coverage Neandertal genome from Vindija Cave in Croatia. Science 358, 655–658 (2017).

4. Mafessoni, F. et al. A high-coverage Neandertal genome from Chagyrskaya Cave. Proc. Natl. Acad. Sci. U.S.A. 117, 15132–15136 (2020).

5. Pääbo, S. The Human Condition—A Molecular Approach. Cell 157, 216–226 (2014).

6. Caporale, N. et al. Tile by tile: capturing the evolutionary mosaic of human conditions. Current Opinion in Genetics & Development 90, 102297 (2025).

7. Zeberg, H., Jakobsson, M. & Pääbo, S. The genetic changes that shaped Neandertals, Denisovans, and modern humans. Cell 187, 1047–1058 (2024).

8. Neubauer, S., Hublin, J.-J. & Gunz, P. The evolution of modern human brain shape. Sci. Adv. 4, eaao5961 (2018).

9. Kuhlwilm, M. & Boeckx, C. A catalog of single nucleotide changes distinguishing modern humans from archaic hominins. Sci Rep 9, 8463 (2019).

10. Cheroni, C. et al. Benchmarking brain organoid recapitulation of fetal corticogenesis. Transl Psychiatry 12, 520 (2022).

11. Trujillo, C. A. et al. Reintroduction of the archaic variant of *NOVA1* in cortical organoids alters neurodevelopment. Science 371, eaax2537 (2021).

12. Mora-Bermúdez, F. et al. Longer metaphase and fewer chromosome segregation errors in modern human than Neanderthal brain development. Sci. Adv. 8, eabn7702 (2022).

13. Pinson, A. et al. Human TKTL1 implies greater neurogenesis in frontal neocortex of modern humans than Neanderthals. Science 377, eabl6422 (2022).

14. Weiss, C. V. et al. The cis-regulatory effects of modern human-specific variants. eLife 10, e63713 (2021).

15. Dutrow, E. V. et al. Modeling uniquely human gene regulatory function via targeted humanization of the mouse genome. Nat Commun 13, 304 (2022).

16. Leonardi, O. et al. CHD2 Dosage Ties Autolysosomal Pathway to Cortical Maturation in Disease and Evolution. bioRxiv Preprint at 10.1101/2025.01.21.634145 (2025).

17. Boettcher, M. et al. Dual gene activation and knockout screen reveals directional dependencies in genetic networks. Nat Biotechnol 36, 170–178 (2018).

18. Najm, F. J. et al. Orthologous CRISPR–Cas9 enzymes for combinatorial genetic screens. Nat Biotechnol 36, 179–189 (2018).

19. Moriano, J. & Boeckx, C. Modern human changes in regulatory regions implicated in cortical development. BMC Genomics 21, 304 (2020).

20. Moriano, J., Leonardi, O., Vitriolo, A., Testa, G. & Boeckx, C. A multi-layered integrative analysis reveals a cholesterol metabolic program in outer radial glia with implications for human brain evolution. Development 151, dev202390 (2024).

21. Bhaduri, A. et al. An atlas of cortical arealization identifies dynamic molecular signatures. Nature 598, 200–204 (2021).

22. Caporale, N. et al. Multiplexing cortical brain organoids for the longitudinal dissection of developmental traits at single-cell resolution. Nat Methods 22, 358–370 (2025).

23. Pereira, M. F. et al. YY1 mutations disrupt corticogenesis through a cell type specific rewiring of cell-autonomous and non-cell-autonomous transcriptional programs. Mol Psychiatry (2025)

24. Peyrégne, S., Boyle, M. J., Dannemann, M. & Prüfer, K. Detecting ancient positive selection in humans using extended lineage sorting. Genome Res. 27, 1563–1572 (2017).

25. Vernot, B. et al. Excavating Neandertal and Denisovan DNA from the genomes of Melanesian individuals. Science 352, 235–239 (2016).

26. Sankararaman, S., Mallick, S., Patterson, N. & Reich, D. The Combined Landscape of Denisovan and Neanderthal Ancestry in Present-Day Humans. Current Biology 26, 1241– 1247 (2016).

27. Kuderna, L. F. K. et al. A global catalog of whole-genome diversity from 233 primate species. Science 380, 906–913 (2023).

28. Yoon, S.-J. et al. Reliability of human cortical organoid generation. Nat Methods 16, 75– 78 (2019).

29. He, Z. et al. An integrated transcriptomic cell atlas of human neural organoids. Nature 635, 690–698 (2024).

30. Pașca, S. P. et al. A framework for neural organoids, assembloids and transplantation studies. Nature (2024)

31. Dixit, A. et al. Perturb-Seq: Dissecting Molecular Circuits with Scalable Single-Cell RNA Profiling of Pooled Genetic Screens. Cell 167, 1853–1866.e17 (2016).

32. Chung, C., Girgiss, J. & Gleeson, J. G. A comparative view of human and mouse telencephalon inhibitory neuron development. Development 152, dev204306 (2025).

33. Delgado, R. N. et al. Individual human cortical progenitors can produce excitatory and inhibitory neurons. Nature 601, 397–403 (2022).

34. Chen, S.-W. et al. Efficient conversion of human induced pluripotent stem cells into microglia by defined transcription factors. Stem Cell Reports 16, 1363–1380 (2021).

35. Hsiung, C. C.-S. et al. Engineered CRISPR-Cas12a for higher-order combinatorial chromatin perturbations. Nat Biotechnol (2024)

36. Liao, C. et al. Convergent coexpression of autism-associated genes suggests some novel risk genes may not be detectable in large-scale genetic studies. Cell Genomics 3, 100277 (2023).

37. Jin, X. et al. In vivo Perturb-Seq reveals neuronal and glial abnormalities associated with autism risk genes. Science 370, eaaz6063 (2020).

38. Schrode, N. et al. Synergistic effects of common schizophrenia risk variants. Nat Genet 51, 1475–1485 (2019).

39. Dräger, N. M. et al. A CRISPRi/a platform in human iPSC-derived microglia uncovers regulators of disease states. Nat Neurosci 25, 1149–1162 (2022).

40. Tian, R. et al. Genome-wide CRISPRi/a screens in human neurons link lysosomal failure to ferroptosis. Nat Neurosci 24, 1020–1034 (2021).

41. Li, C. et al. Single-cell brain organoid screening identifies developmental defects in autism. Nature 621, 373–380 (2023).

42. Gokhman, D. et al. Differential DNA methylation of vocal and facial anatomy genes in modern humans. Nat Commun 11, 1189 (2020).

43. Dutto, I. et al. Pathway-specific effects of ADSL deficiency on neurodevelopment. eLife 11, e70518 (2022).

44. Wang, L. et al. ADSL promotes autophagy and tumor growth through fumarate-mediated Beclin1 dimethylation. Nat Chem Biol (2025) doi:10.1038/s41589-024-01825-9.

45. Pravata, M. Veronica et al. DCHS1 Modulates Forebrain Proportions in Modern Humans via a Glycosylation Change.

46. Molz, B. et al. Imaging genomics reveals genetic architecture of the globular human braincase. BiorXiv Preprint at 10.1101/2024.03.20.585712 (2024).

47. Stepanova, V. et al. Reduced purine biosynthesis in humans after their divergence from Neandertals. eLife 10, e58741 (2021).

48. Panagiotakos, G. & Pasca, S. P. A matter of space and time: Emerging roles of disease-associated proteins in neural development. Neuron 110, 195–208 (2022).

49. Birtele, M. et al. Non-synaptic function of the autism spectrum disorder-associated gene SYNGAP1 in cortical neurogenesis. Nat Neurosci 26, 2090–2103 (2023).

50. Paten, B., Herrero, J., Beal, K., Fitzgerald, S. & Birney, E. Enredo and Pecan: Genome-wide mammalian consistency-based multiple alignment with paralogs. Genome Res. 18, 1814–1828 (2008).

51. Yan, G. et al. Genome sequencing and comparison of two nonhuman primate animal models, the cynomolgus and Chinese rhesus macaques. Nat Biotechnol 29, 1019–1023 (2011).

52. Sherry, S. T. dbSNP: the NCBI database of genetic variation. Nucleic Acids Research 29, 308–311 (2001).

53. Sloan, S. A., Andersen, J., Pașca, A. M., Birey, F. & Pașca, S. P. Generation and assembly of human brain region–specific three-dimensional cultures. Nat Protoc 13, 2062–2085 (2018).

54. Doench, J. G. et al. Optimized sgRNA design to maximize activity and minimize off-target effects of CRISPR-Cas9. Nat Biotechnol 34, 184–191 (2016).

55. Sanson, K. R. et al. Optimized libraries for CRISPR-Cas9 genetic screens with multiple modalities. Nat Commun 9, 5416 (2018).

56. Chavez, A. et al. Highly efficient Cas9-mediated transcriptional programming. Nat Methods 12, 326–328 (2015).

57. Korsunsky, I. et al. Fast, sensitive and accurate integration of single-cell data with Harmony. Nat Methods 16, 1289–1296 (2019).

58. Polioudakis, D. et al. A Single-Cell Transcriptomic Atlas of Human Neocortical Development during Mid-gestation. Neuron 103, 785–801.e8 (2019).

59. Zanella, M. et al. Dosage analysis of the 7q11.23 Williams region identifies *BAZ1B* as a major human gene patterning the modern human face and underlying self-domestication. Sci. Adv. 5, eaaw7908 (2019).

60. Papalexi, E. et al. Characterizing the molecular regulation of inhibitory immune checkpoints with multimodal single-cell screens. Nat Genet 53, 322–331 (2021).

61. Kamimoto, K. et al. Dissecting cell identity via network inference and in silico gene perturbation. Nature 614, 742–751 (2023).

62. Nakamura, T. et al. Topologically associating domains define the impact of de novo promoter variants on autism spectrum disorder risk. Cell Genomics 4, 100488 (2024).

63. Gandal, M. J. et al. Transcriptome-wide isoform-level dysregulation in ASD, schizophrenia, and bipolar disorder. Science 362, eaat8127 (2018).

64. De La Torre-Ubieta, L. et al. The Dynamic Landscape of Open Chromatin during Human Cortical Neurogenesis. Cell 172, 289–304.e18 (2018).

