## Supplementary Materials for "Regulatory logic of human cortex evolution by combinatorial perturbations"

**The PDF file includes:**

Figs. S1 to S10
Tables S1 to S2

**Fig. S1.**

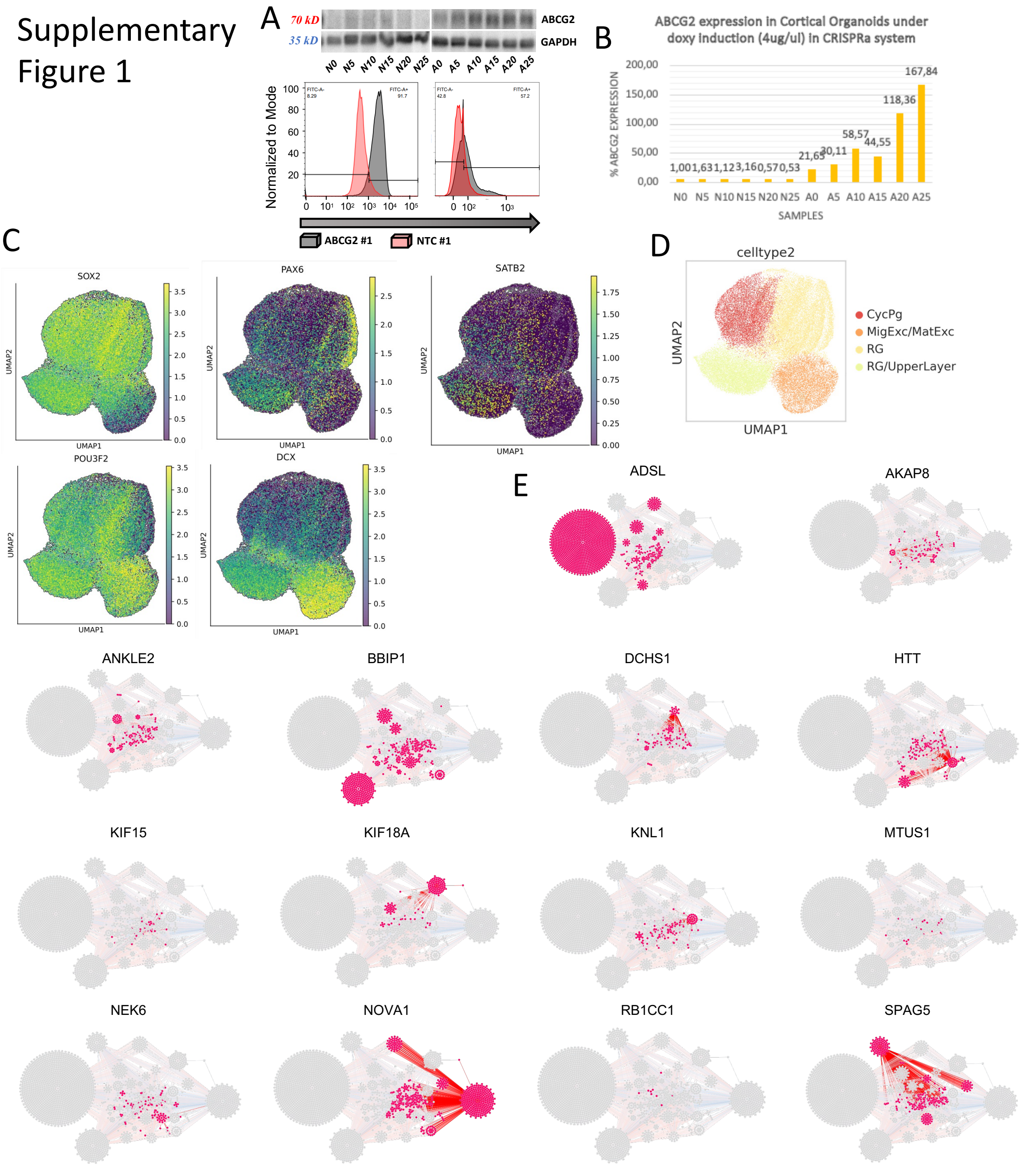


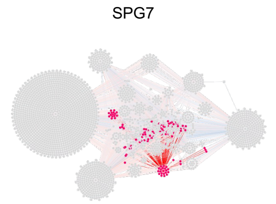


Supplementary Figure 1 Caption

A) Western blot of ABCG2 and GAPDH coupled with Fluorescein isothiocyanate cytometry (FITC-A) mean intensity upon doxycycline CRISPRa induction at different time points show stable up-regulation of ABCG2; B) Barplot of percentage (%) of gene expression in induced CRISPRa line vs non induced lines at 4ug/uL doxycycline concentration; C) Cortical brain organoids at 25 DIV show broad expression (UMI normalized counts) of SOX2 and PAX6 in cells identified as cycling progenitors (CycPg) and radial glia (RG), POU3F2 a marker of migratory neurons is expressed in a small cluster of SATB2$^{+}$ cells that we called RG/UpperLayer and double POU3F2$^{+}$/DCX$^{+}$ cells were annotated as migratory/maturing excitatory neurons (MigExc/MatExc); D) Cell type annotation is reported next to markers expression UMAPs. E) Genes co-expressed with our candidate genes across cell types in 25 DIV CBO are indicated in pink for each candidate, on top of the full co-expression network.

**Fig. S2.**

Supplementary Figure 2

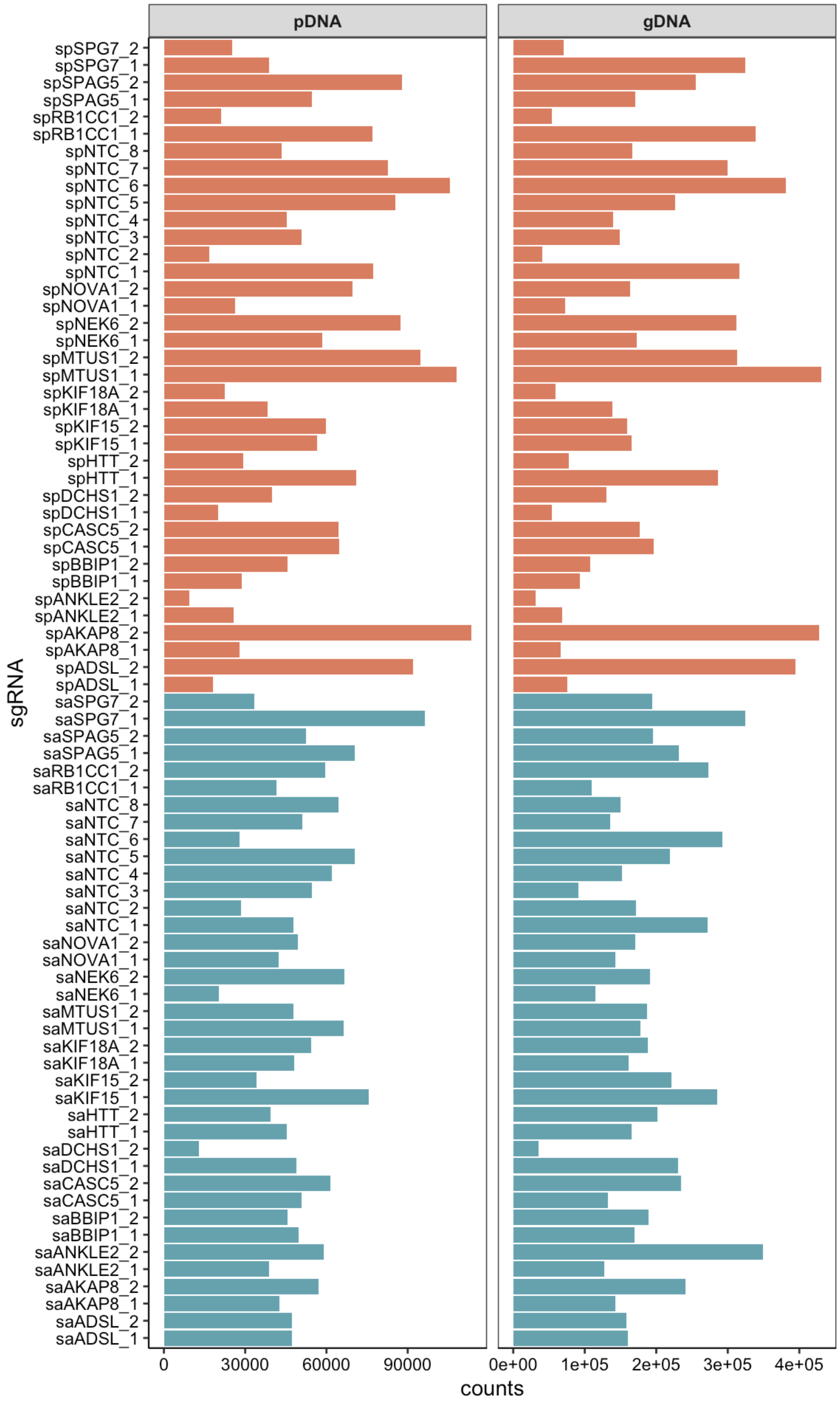


Supplementary Figure 2. Plasmid and Genomic DNA comparison. Bar plot reporting the total sequenced counts (x-axis) for each potential sgRNA (y-axis). Guides designed for upregulation are reported in light red and guides designed for downregulation are reported in light blue. NTC guides are also reported.

**Fig. S3.**

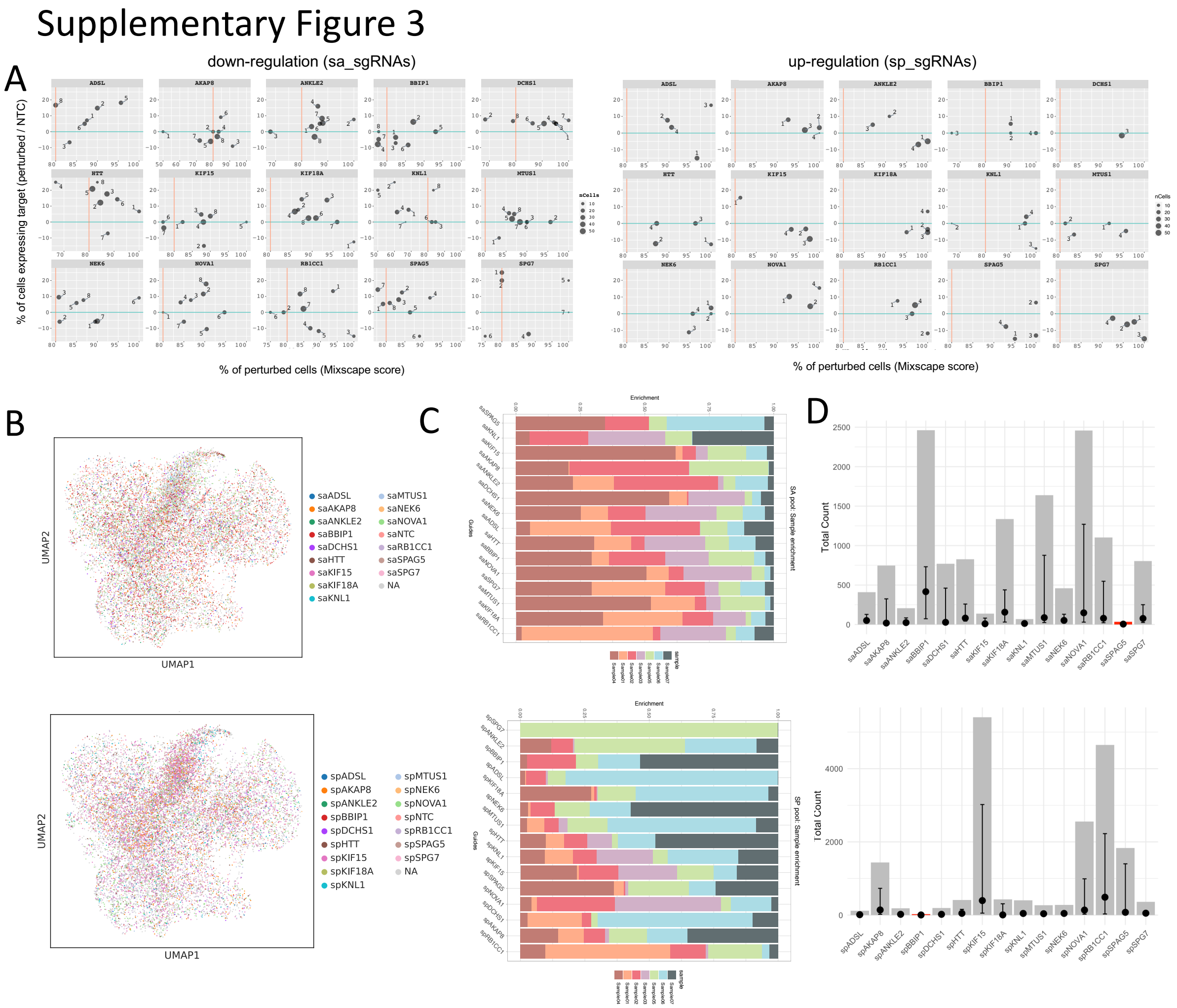


Supplementary Figure 3. Guide selection and detection quality controls

A) Scatter plot reporting the detection rate of the perturbed gene (y-axis) in cells bearing a certain single perturbation sgRNA candidate guide, with respect to the detection rate of the same gene in NTC cells (down-regulation on the left, and up-regulation on the right) compared to Mixscape score (x-axis, percentage of perturbed cells among those bearing the same sgRNA). The data is derived from single-perturbation iPSC mosaic coltures; B) UMAP of mosaic CBOs showing distribution of cells bearing sgRNA guides for downregulation (above), and for upregulation (below); C) Stacked bar plots reporting percentage of abundance of each perturbation in CBO cells (x-axis). Different colors represent the seven different sequencing runs. Each individual perturbation (downregulation above, upregulations below) is reported on the y-axis; D) Bar plots of total cell counts per perturbation across CBO cells (grey), summing all seven sequencing runs. A dot is reported for the median number of cells with each perturbation across sequencing runs. Highest value is reported with a black internal bar (black).

**Fig. S4.**

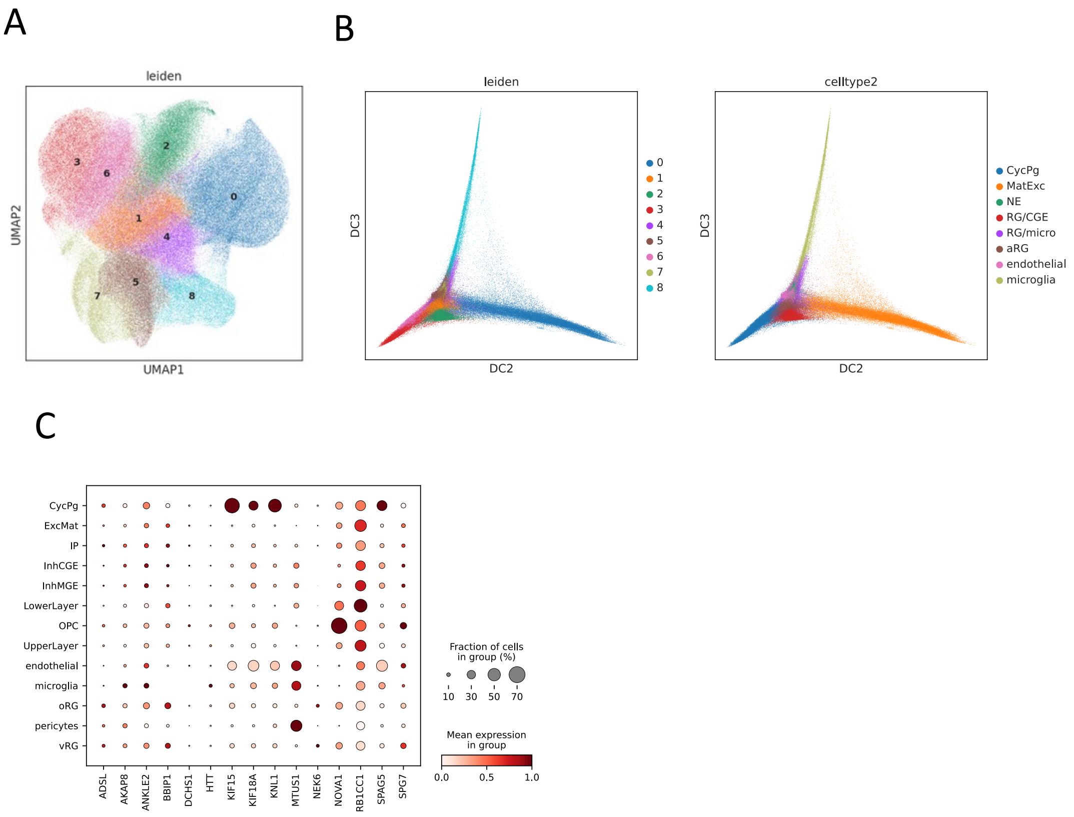


Supplementary Figure 4. Cell type characterization

A) UMAP of CBOs data (n = 210,257). Leiden clustering is reported by colouring cells, and cluster numbers reported within each group of cells. B) Diffusion maps are reported for the same data in panel A, with matching colors for leiden clusters (left panel) and cell type names reported on the right panel; C) Expression levels of our core genes identified in data from Polioudakis et al[58]

**Fig. S5.**

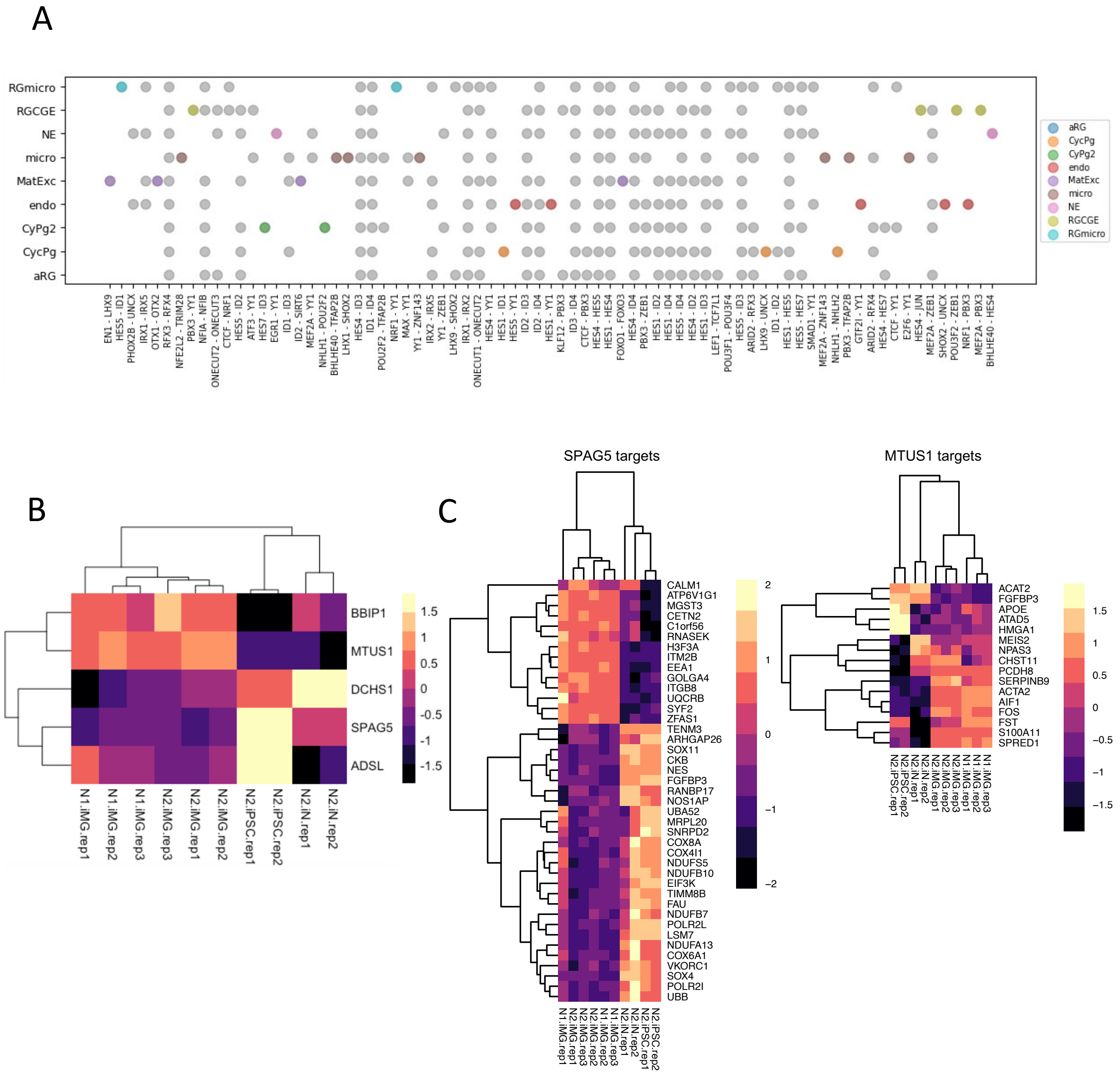


Supplementary Figure 5. Validation of cell type annotation via TF-TF interaction and external data

A) Dot plot representing the genetic interactions between couples of TFs. From each cell type specific GRN generated with CellOracle, an hypergeometic test was performed to identify significant overlaps between lists of TF targets. Grey dots represent interactions between TFs that are not cell type specific, Cell type-specific TF-TF significant overlaps are colored by cell type as per the legend on the right. Names of cell types are also reported on the left (y-axis). Names of TFs involved in each couple is reported on the bottom (x-axis); B) Heatmap reporting z-scores of log(TMM)+1 normalized values calculated from the read counts of GSE163984 bulk RNA-seq data from iPSC (N2.iPSC.rep1 and N2.iPSC.rep2), iPSC-derived neurons (N2.iN.rep1 and N2.iN.rep2), and iPSC-derived microglia (N1.iMG.rep1-3 and N2.iMG.rep1-3). Expression values for the core genes differentially expressed across cell types (ADSL, BBIP1, DCHS1, MTUS1 and SPAG5) are reported; C) Heatmap reporting z-scores of log(TMM)+1 normalized values of SPAG5 (left) and MTUS1 (right) targets differentially expressed across cel types in the same data reported for panel B. Majority of SPAG5 targets are downregulated in microglia (28 out of 42), including neuronal differentiation genes such as NES, SOX4 and SOX11. Majority of MTUS1 targets are upregulated in microglia (11 out of 16).

**Fig. S6.**

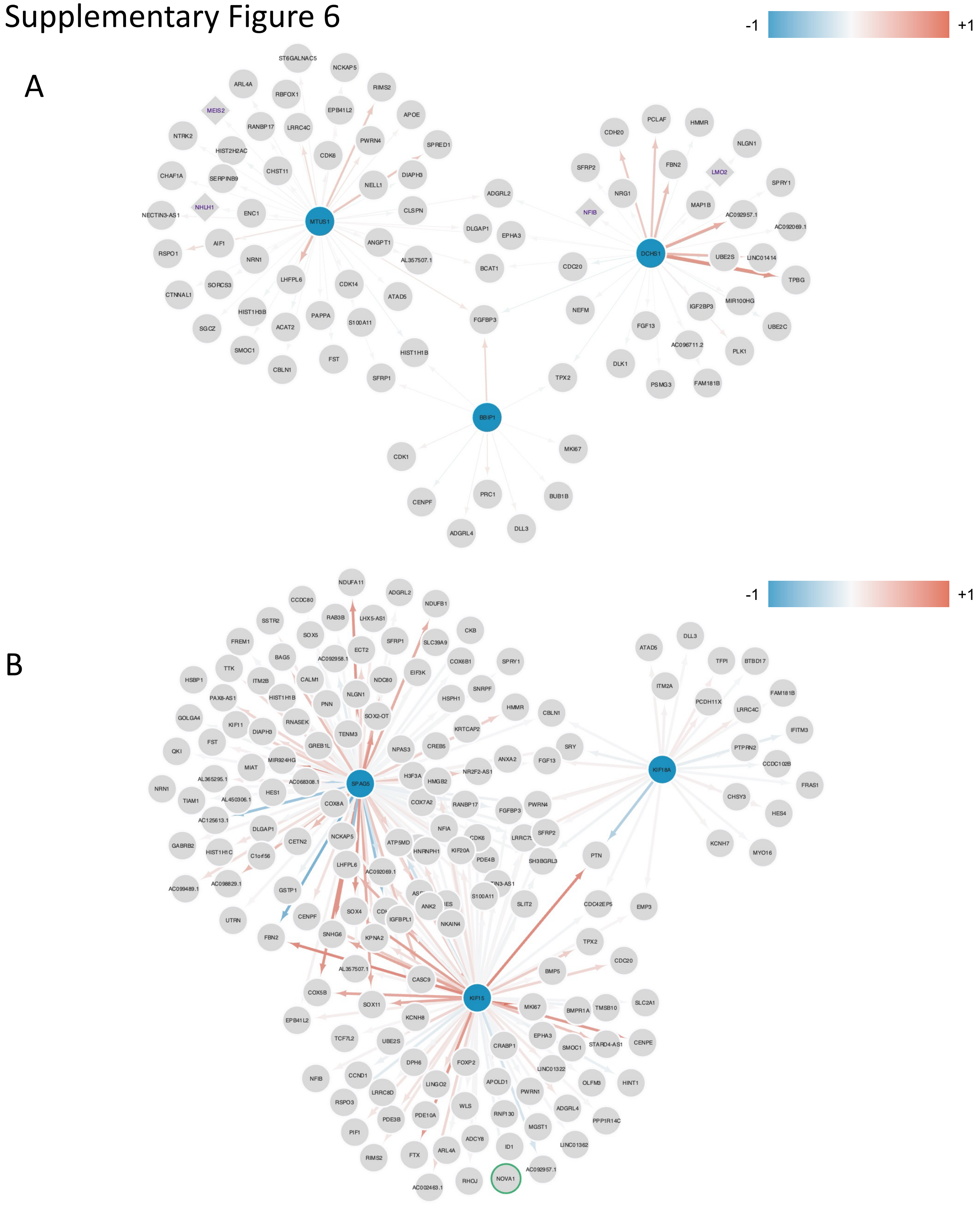


Supplementary Figure 6. Subsets of gene regulatory networks (GRN) involving core genes in microglia-like and RG/CGE. A) Graph of microglia-like cells interactions identified for BBIP1, DCHS1 and MTUS1. Core genes are reported as blue circles (nodes), red arrows (edges) indicate positive expression correlations, and blue arrows represent inverse coexpression. Genes identified as TF in the GRN are reported as square nodes with gene names reported in purple; Graph of RG/CGE cells interactions identified for KIF15, KIF18A and SPAG5. Core genes are reported as blue circles (nodes), red arrows (edges) indicate positive expression correlations, and blue arrows represent inverse coexpression. Other core genes are circled in green (NOVA1 here, target of KIF15).

**Fig. S7.**

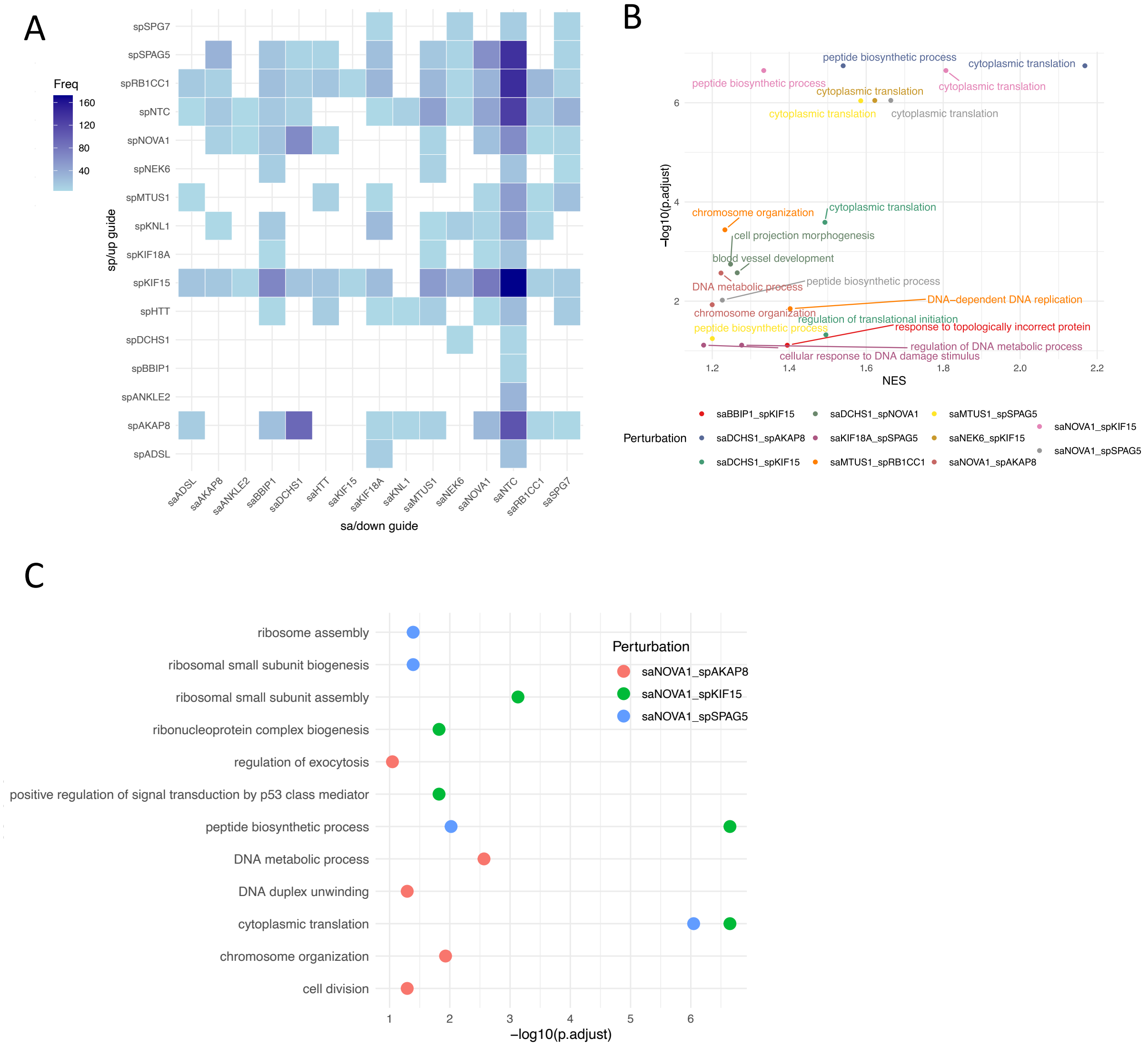


Supplementary Figure 7
A) Number of cells ( see “Freq” color bar) found bearing each of the indicated couple of downregulation (x-axis) and upregulation (y-axis) guides; B) Scatter plot of top Gene Ontology categories enriched in *biological processes* for all couples of perturbations that were found in more than 40 cells and showed differentially expressed genes (FDR < 0.05 and |FC| > 1.5). Perturbation couples are indicated by color. Normalized enrichment scores from gene set enrichment analysis (GSEA) is reported on the x-axis, and significance (-log10 adjusted p-value) is reported on the y-axis; C) Scatter plot of top Gene Ontology enrichments obtained by topGO for the three double perturbations involving NOVA1. Dots are colored by perturbation, x-axis reports significance (-log10 p-adjusted), y-axis report GO categories.

**Fig. S8.**

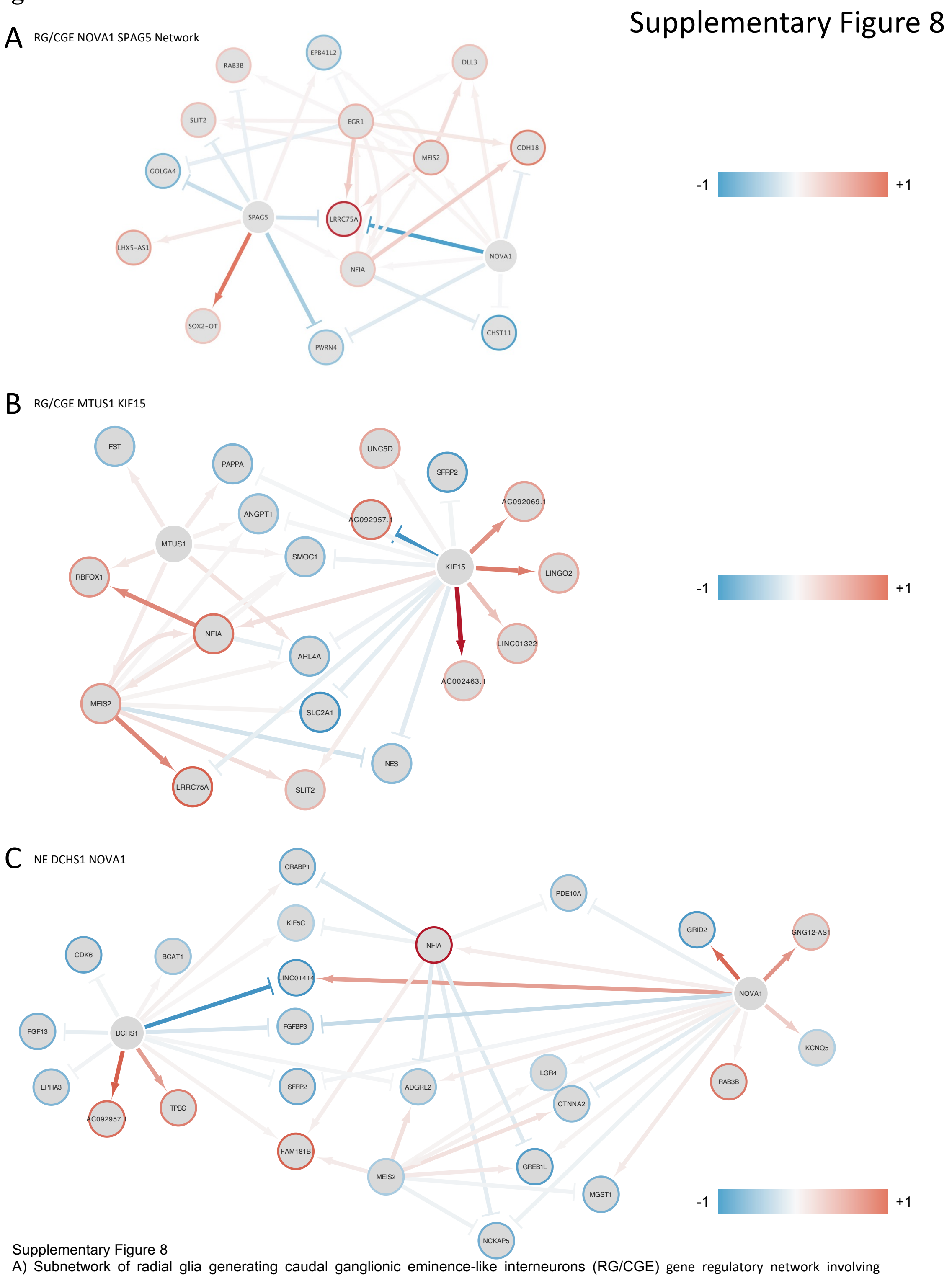


Supplementary Figure 8

A) Subnetwork of radial glia generating caudal ganglionic eminence-like interneurons (RG/CGE) gene regulatory network involving NOVA1 and SPAG5 cell type specific unique and shared targets.

B) Subnetwork of RG/CGE gene regulatory network involving KIF15 and MTUS1 cell type specific unique and shared targets.

C) Subnetwork of neural epithelium (NE) gene regulatory network involving DCHS1 and NOVA1 cell type specific unique and shared targets.

All networks report nodes as circle, colored in blue or red if they were down- or up-regulated in the differential expression analysis of double perturbations. First gene reported in the figure title is down-regulated, second gene reported was up-regulated. Link between nodes (edges) are colored in blue or red if the regulator (core genes) was inversely or directly co-expressed with the target in the specific cell type.

**Fig.S9.**

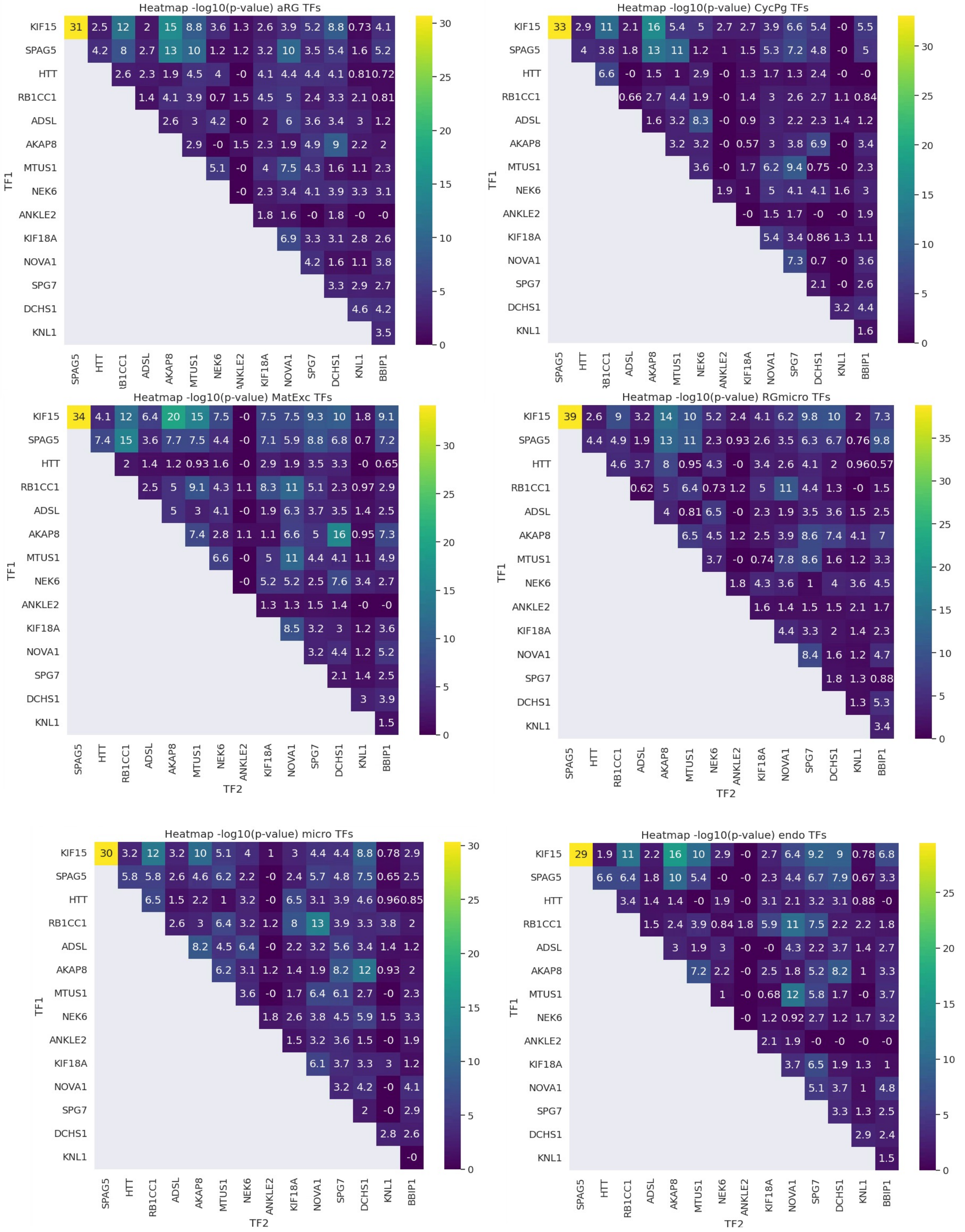


Supplementary Figure 9

Hetmaps reporting for each cell type the significance of the overlap between targets of core genes in each cell type. Significance was measured by hypergeometric test, comparing number of targets shared by each couple (intersection fo x- and y-axis) with respect to the union of targets of all core genes. Cell type are reported on top of each heatmap.

**Fig. S10.**

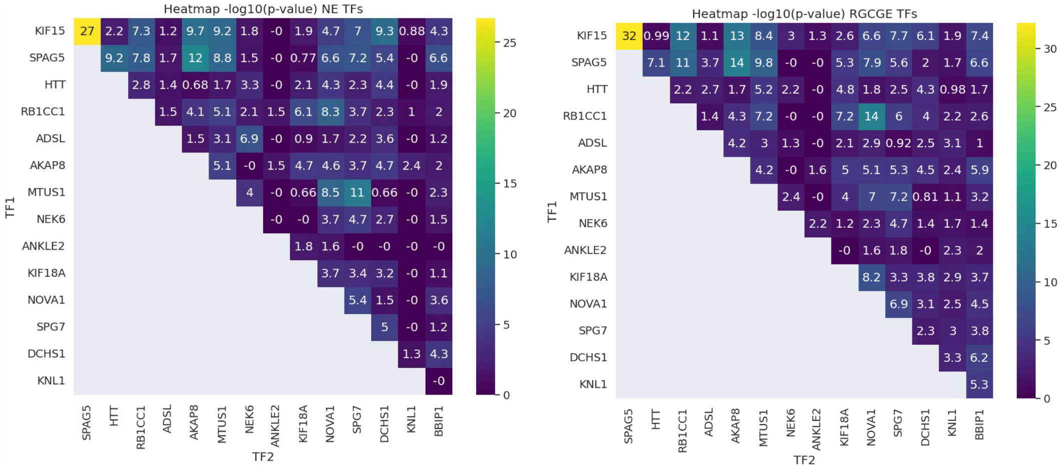


Supplementary Figure 10
Hetmaps reporting for each cell type the significance of the overlap between targets of core genes in each cell type. Significance was measured by hypergeometric test, comparing number of targets shared by each couple (intersection fo x- and y-axis) with respect to the union of targets of all core genes. Cell type are reported on top of each heatmap.

Table S1.

| sgRNA_barcode | counts | sgRNA_id | sgRNA_seq | gene |
|---|---|---|---|---|
| GAAAACGCTCGACAAC | 79432 | SPAG5_1 | GAAAACGCTCGACAACGGAG | SPAG5 |
| CGGAAAACGCTCGACA | 140313 | SPAG5_2 | CGGAAAACGCTCGACAACGG | SPAG5 |
| GGAAAACGCTCGACAA | 140893 | SPAG5_3 | GGAAAACGCTCGACAACGGA | SPAG5 |
| CAGCGGAAAACGCTCG | 97973 | SPAG5_4 | CAGCGGAAAACGCTCGACAA | SPAG5 |
| TGGCGCCAAGAAGCTC | 78317 | KIF18A_1 | TGGCGCCAAGAAGCTCAGCC | KIF18A |
| AGCTCAGCCTGGCTGC | 94422 | KIF18A_2 | AGCTCAGCCTGGCTGCGCAG | KIF18A |
| TCGGACGCTGTCACGG | 71038 | KIF18A_3 | TCGGACGCTGTCACGGCTCG | KIF18A |
| AAGCTCAGCCTGGCTG | 30395 | KIF18A_4 | AAGCTCAGCCTGGCTGCGCA | KIF18A |
| ATAAAATTGGCACGGA | 64590 | CASC5_1 | ATAAAATTGGCACGGAGTCT | CASC5 |
| TCTTAGAAATAGGGTT | 48592 | CASC5_2 | TCTTAGAAATAGGGTTGGTC | CASC5 |
| AGGAGTCTTAGAAATA | 37182 | CASC5_3 | AGGAGTCTTAGAAATAGGGT | CASC5 |
| TGAAGGAAATAAAATT | 41924 | CASC5_4 | TGAAGGAAATAAAATTGGCA | CASC5 |
| GCCTGAGCCCGTCTAC | 86001 | SPG7_1 | GCCTGAGCCCGTCTACCTCG | SPG7 |
| GGGTGCGGCAAGCGGC | 59633 | SPG7_2 | GGGTGCGGCAAGCGGCACCC | SPG7 |
| GTAGACGGGCTCAGGC | 74756 | SPG7_3 | GTAGACGGGCTCAGGCTCGG | SPG7 |
| GAGGTAGACGGGCTCA | 100530 | SPG7_4 | GAGGTAGACGGGCTCAGGCT | SPG7 |
| AACCACTTGAAAGATC | 37215 | MTUS1_1 | AACCACTTGAAAGATCGCTA | MTUS1 |
| TCCCCTAGCGATCTTT | 51947 | MTUS1_2 | TCCCCTAGCGATCTTTCAAG | MTUS1 |
| TAACCACTTGAAAGAT | 79045 | MTUS1_3 | TAACCACTTGAAAGATCGCT | MTUS1 |
| TACAGGATCTGTGACA | 75301 | MTUS1_4 | TACAGGATCTGTGACATTAT | MTUS1 |
| AGCGAACCAAGTCAAT | 76113 | ADSL_1 | AGCGAACCAAGTCAATTCTC | ADSL |
| CAGCGGAGATTCTCCG | 81978 | ADSL_2 | CAGCGGAGATTCTCCGCAGC | ADSL |
| CTTCTGCCTGAGAATT | 98772 | ADSL_3 | CTTCTGCCTGAGAATTGACT | ADSL |
| TCCGCAGCCGGCAGCG | 91549 | ADSL_4 | TCCGCAGCCGGCAGCGCCCA | ADSL |
| GACGGCGGCCCCGCTC | 124288 | ANKLE2_1 | GACGGCGGCCCCGCTCGCGG | ANKLE2 |
| CCGCTTGGGTCCCAGG | 57124 | ANKLE2_2 | CCGCTTGGGTCCCAGGGCCG | ANKLE2 |
| GCGCCGCGCTCGAGGC | 67752 | ANKLE2_3 | GCGCCGCGCTCGAGGCGGCC | ANKLE2 |
| GGCGCCGCGCTCGAGG | 129042 | ANKLE2_4 | GGCGCCGCGCTCGAGGCGGC | ANKLE2 |
| TCAACAGTAACTTCCG | 80152 | BBIP1_1 | TCAACAGTAACTTCCGATCG | BBIP1 |
| CTTCAACAGTAACTTC | 88158 | BBIP1_2 | CTTCAACAGTAACTTCCGAT | BBIP1 |
| TACAGAGTAAGCCTCC | 73079 | BBIP1_3 | TACAGAGTAAGCCTCCAGCA | BBIP1 |
| TTGTCACTCTACCCTG | 85074 | BBIP1_4 | TTGTCACTCTACCCTGCTGG | BBIP1 |
| GTCTACGCAGGTCCGG | 36873 | RB1CC1_1 | GTCTACGCAGGTCCGGAAGT | RB1CC1 |
| CCAAACACTGGATCGT | 80919 | RB1CC1_2 | CCAAACACTGGATCGTCCCC | RB1CC1 |
| GAGAGGCGTCTACGCA | 91283 | RB1CC1_3 | GAGAGGCGTCTACGCAGGTC | RB1CC1 |
| GGCCAGGACAGCCAAC | 38263 | RB1CC1_4 | GGCCAGGACAGCCAACCTGC | RB1CC1 |
| TTGCCCGGCCTACGCA | 53781 | KIF15_1 | TTGCCCGGCCTACGCATCGG | KIF15 |
| ACGGATTCGCGACGAT | 90494 | KIF15_2 | ACGGATTCGCGACGATACAG | KIF15 |
| GCGTAGGCCGGGCAAC | 102978 | KIF15_3 | GCGTAGGCCGGGCAACATGG | KIF15 |
| TCTGGCCCGCTTCAAG | 41042 | KIF15_4 | TCTGGCCCGCTTCAAGGAAC | KIF15 |
| GGACCCGAGCGCGAAG | 102501 | NOVA1_1 | GGACCCGAGCGCGAAGTGTG | NOVA1 |
| GACCCGAGCGCGAAGT | 57312 | NOVA1_2 | GACCCGAGCGCGAAGTGTGT | NOVA1 |
| CTGTGAGGGGAGGGTA | 38561 | NOVA1_3 | CTGTGAGGGGAGGGTAGAGA | NOVA1 |
| TCCCGCTTGTTGGGGG | 63235 | NOVA1_4 | TCCCGCTTGTTGGGGGGGGC | NOVA1 |
| TGCCGCCACGCAAGCT | 45716 | NEK6_1 | TGCCGCCACGCAAGCTTGAG | NEK6 |
| CGCCCCTCAAGCTTGC | 58877 | NEK6_2 | CGCCCCTCAAGCTTGCGTGG | NEK6 |
| CAGAACTCGGCGGAGT | 53265 | NEK6_3 | CAGAACTCGGCGGAGTCGAG | NEK6 |
| GCAGAACTCGGCGGAG | 113169 | NEK6_4 | GCAGAACTCGGCGGAGTCGA | NEK6 |
| CCATTGGCTGGTCTTC | 43431 | AKAP8_1 | CCATTGGCTGGTCTTCACGA | AKAP8 |
| CTGGATCGTAGCAGCC | 56329 | AKAP8_2 | CTGGATCGTAGCAGCCCGAA | AKAP8 |
| GCTGCTACGATCCAGC | 37212 | AKAP8_3 | GCTGCTACGATCCAGCTCCG | AKAP8 |
| CCACAACTCCCATTAG | 54192 | AKAP8_4 | CCACAACTCCCATTAGGCCA | AKAP8 |
| CCTTCCCCCCGGCGCC | 56812 | DCHS1_1 | CCTTCCCCCCGGCGCCCGCC | DCHS1 |
| CCGCCGCCCCCCCTTC | 62413 | DCHS1_2 | CCGCCGCCCCCCCTTCCCCC | DCHS1 |
| GGGGCGGCGGCGCGGA | 78289 | DCHS1_3 | GGGGCGGCGGCGCGGAGCGC | DCHS1 |
| AGGGGCGGAGGGAGGC | 39369 | DCHS1_4 | AGGGGCGGAGGGAGGCCGGG | DCHS1 |
| TGATTGACAGCCCTAG | 69044 | HTT_1 | TGATTGACAGCCCTAGCCTG | HTT |
| GGGCTGTCAATCATGC | 78776 | HTT_2 | GGGCTGTCAATCATGCTGGC | HTT |
| TTGGCAGAGTCCGCAG | 87825 | HTT_3 | TTGGCAGAGTCCGCAGGCTA | HTT |
| GTCAATCATGCTGGCC | 78357 | HTT_4 | GTCAATCATGCTGGCCGGCG | HTT |

**Supplementary Table 1**: Genomic sequencing raw read counts from pre-screening organoids for spCas9 and saCas9 respectively

| sgRNA_seq | counts | sgRNA_id | gene | plotcolor |
|---|---|---|---|---|
| AGAACCTTCGGCGCTCTGACA | 3464 | SPAG5_1 | SPAG5 | #65a2ae |
| TGTCTCCGCACGCCGAAGTGA | 7376 | SPAG5_2 | SPAG5 | #65a2ae |
| GGAAGGTCCTGTCATTAGGCA | 4939 | SPAG5_3 | SPAG5 | #65a2ae |
| AAGATCCTAGAACAGATAGAC | 8236 | SPAG5_4 | SPAG5 | #65a2ae |
| AAGTGACAAGAGTGCTTTCAC | 11283 | SPAG5_5 | SPAG5 | #65a2ae |
| GTCACTTCCAAGCAAGGGCAC | 6953 | SPAG5_6 | SPAG5 | #65a2ae |
| AGTCGGTCCCGACAACTCACA | 8936 | SPAG5_7 | SPAG5 | #65a2ae |
| TTTAGGAGTAGAGCTAATTTG | 57302 | SPAG5_8 | SPAG5 | #65a2ae |
| CTTTGCCCGGTTAGCATACTT | 20172 | KIF18A_1 | KIF18A | #65a2ae |
| ATCAGCTGATCCTAGCATAGT | 14723 | KIF18A_2 | KIF18A | #65a2ae |
| ACCAAAGCAAAACGATCTACC | 7496 | KIF18A_3 | KIF18A | #65a2ae |
| AGAGTGTTATATGTGTCATCG | 12451 | KIF18A_4 | KIF18A | #65a2ae |
| GTCGAACACGTTTGGCAAATC | 73450 | KIF18A_5 | KIF18A | #65a2ae |
| ATATCCATTCAAAAAACTACG | 14166 | KIF18A_6 | KIF18A | #65a2ae |
| ACAGGCATAGAATGCTTAGTT | 17617 | KIF18A_7 | KIF18A | #65a2ae |
| AGAATCGAGAAGAAATTAGAC | 3880 | KIF18A_8 | KIF18A | #65a2ae |
| CATCCAGTGAGACCAACTCAC | 15292 | CASC5_1 | CASC5 | #65a2ae |
| GCAGCTTATAATCTAGCACCG | 2065 | CASC5_2 | CASC5 | #65a2ae |
| GCTTGTCTGAGTGGGATGTCG | 7930 | CASC5_3 | CASC5 | #65a2ae |
| AGAAGATCGATAACTGCCTCA | 24415 | CASC5_4 | CASC5 | #65a2ae |
| CTGCAATGGTGCACTACCAGT | 215931 | CASC5_5 | CASC5 | #65a2ae |
| CCAGAATATTAGCCATGACCC | 6782 | CASC5_6 | CASC5 | #65a2ae |
| GTGATATGTACTCCTACTGAG | 842 | CASC5_7 | CASC5 | #65a2ae |
| ATTATTAGACCATTCCTCCAG | 1930 | CASC5_8 | CASC5 | #65a2ae |
| AGTCAGCTTCAAAGACGTGGC | 4404 | SPG7_1 | SPG7 | #65a2ae |
| GCCCAGGGCGGCGTTTGTCCG | 22454 | SPG7_2 | SPG7 | #65a2ae |
| CTCCACGGCGTACTCGAAGTT | 23193 | SPG7_3 | SPG7 | #65a2ae |
| CTCATCGATGTAGACGATGCA | 6605 | SPG7_4 | SPG7 | #65a2ae |
| CTTCCGTGCACACTCTCAACT | 16682 | SPG7_5 | SPG7 | #65a2ae |
| CTTCACCATGGAGTAGGCGAT | 21704 | SPG7_6 | SPG7 | #65a2ae |
| AAACTGGAAGTCCGCGAGTTT | 19220 | SPG7_7 | SPG7 | #65a2ae |
| AGAGGAGAGGAGACGCCGTGA | 0 | SPG7_8 | SPG7 | #65a2ae |
| CAGAACGAGAGAATCGGCTTA | 10293 | MTUS1_1 | MTUS1 | #65a2ae |
| TCCTTTGGCCGTCAGTCGTAT | 18160 | MTUS1_2 | MTUS1 | #65a2ae |
| GAGAAGCAGGAATCGCTAGAG | 12354 | MTUS1_3 | MTUS1 | #65a2ae |
| TTCACATACGACTGACGGCCA | 16694 | MTUS1_4 | MTUS1 | #65a2ae |
| GCCGTCAGTCGTATGTGAATG | 38896 | MTUS1_5 | MTUS1 | #65a2ae |
| GGCTTAAAGAGTTTTACACCA | 8235 | MTUS1_6 | MTUS1 | #65a2ae |
| GTGGCTGGATACATAAGCTTC | 8464 | MTUS1_7 | MTUS1 | #65a2ae |
| CAGGCTGAAAAAACAGAACGA | 4382 | MTUS1_8 | MTUS1 | #65a2ae |
| TGTCTGTAGCGGGTCCATGAC | 8773 | ADSL_1 | ADSL | #65a2ae |
| CCGAGATGACCTGCGCTTCCG | 52011 | ADSL_2 | ADSL | #65a2ae |
| GATCCAGTAAATGATCCAACT | 5961 | ADSL_3 | ADSL | #65a2ae |
| TTTCTTACCGCAGATACTATA | 9130 | ADSL_4 | ADSL | #65a2ae |
| TCAAGTGCGATGCCATATAAG | 14801 | ADSL_5 | ADSL | #65a2ae |
| GACCTGCGCTTCCGGGGAGTA | 5779 | ADSL_6 | ADSL | #65a2ae |
| TCTCGGACACGCTTCAAGTTC | 0 | ADSL_7 | ADSL | #65a2ae |
| GTGAAAGCAGAATTATGTCTG | 2382 | ADSL_8 | ADSL | #65a2ae |
| GAGGTACAGGTCCACCACGTA | 10309 | ANKLE2_1 | ANKLE2 | #65a2ae |
| CTGTTGGCTCGATTGAAACTT | 2870 | ANKLE2_2 | ANKLE2 | #65a2ae |
| TGTGCACGGCCGGGAACTGAT | 15149 | ANKLE2_3 | ANKLE2 | #65a2ae |
| GAAGCCTGCTTTCTCTCGAGG | 4617 | ANKLE2_4 | ANKLE2 | #65a2ae |
| CTAGAAACGCCCTCACGGAAA | 70665 | ANKLE2_5 | ANKLE2 | #65a2ae |
| GCCAGGATCGCTAAAATGTCC | 6750 | ANKLE2_6 | ANKLE2 | #65a2ae |
| CACCTGTCTCGGATTTGACTG | 11414 | ANKLE2_7 | ANKLE2 | #65a2ae |
| GAGGAGCCATCAAAACTCGAT | 19834 | ANKLE2_8 | ANKLE2 | #65a2ae |
| TCATGGCGTTCGTCTAGGATG | 59301 | RB1CC1_1 | RB1CC1 | #65a2ae |
| AGAATGCATGGCTGCAGATCG | 4320 | RB1CC1_2 | RB1CC1 | #65a2ae |
| CACTTCAGGTTAGGCACATAC | 8758 | RB1CC1_3 | RB1CC1 | #65a2ae |
| TAACGTTACTTACCCTGCTCA | 12404 | RB1CC1_4 | RB1CC1 | #65a2ae |
| TCTGGTTAGGCACTCCAACAG | 3114 | RB1CC1_5 | RB1CC1 | #65a2ae |
| CCTCGTAATAGAGCTGTTAGA | 10463 | RB1CC1_6 | RB1CC1 | #65a2ae |
| GCGATAGACAGTAGACGAATG | 30141 | RB1CC1_7 | RB1CC1 | #65a2ae |
| GTAAACTTGACGGACTAATAA | 30658 | RB1CC1_8 | RB1CC1 | #65a2ae |
| GTCACAGATAGAGCACCACCC | 2496 | KIF15_1 | KIF15 | #65a2ae |
| AAGTCGAACGAACCCAAACTT | 16249 | KIF15_2 | KIF15 | #65a2ae |
| GAAAAGTGAGGTACATGACCT | 1350 | KIF15_3 | KIF15 | #65a2ae |
| CAACTGTGGCTAAAAGCATTG | 9815 | KIF15_4 | KIF15 | #65a2ae |
| CTTCTTTAGTCGCACTACGTA | 3927 | KIF15_5 | KIF15 | #65a2ae |
| TTCAGCTCTGTCTGAATTTGC | 8307 | KIF15_6 | KIF15 | #65a2ae |
| AACTCCTGGAAACACAAGAAC | 33941 | KIF15_7 | KIF15 | #65a2ae |
| GTGATTACAGCACTTGTCGAC | 13714 | KIF15_8 | KIF15 | #65a2ae |
| CCAAGAGGACCAATACGGGCG | 4329 | NOVA1_1 | NOVA1 | #65a2ae |
| AATCCAGATCGCATCAAACAA | 7085 | NOVA1_2 | NOVA1 | #65a2ae |
| CAGGTCTATGGGAACCCCAGT | 7202 | NOVA1_3 | NOVA1 | #65a2ae |
| GAAAAGTCCACAGATGGATCC | 43956 | NOVA1_4 | NOVA1 | #65a2ae |
| GGAAAACATTAGTGGAATACC | 3554 | NOVA1_5 | NOVA1 | #65a2ae |
| TGATGGGATCAACTTGCAAGA | 16952 | NOVA1_6 | NOVA1 | #65a2ae |
| TTGAAGCACTGAATGCAGTTC | 2572 | NOVA1_7 | NOVA1 | #65a2ae |
| GGATACAGATCTCCAAAAAAG | 848 | NOVA1_8 | NOVA1 | #65a2ae |
| TCAGCTCGTTGTCTTCGATAA | 76115 | NEK6_1 | NEK6 | #65a2ae |
| TTCTCGGAGTAGTGCTCCCCG | 14726 | NEK6_2 | NEK6 | #65a2ae |
| CAGACAGGTCCCCCTTACCTA | 9493 | NEK6_3 | NEK6 | #65a2ae |
| TTTCGATCTGGAAGTCCGCCA | 31081 | NEK6_4 | NEK6 | #65a2ae |
| GCCCTACTACATGTCACCGGA | 17187 | NEK6_5 | NEK6 | #65a2ae |
| ATCCGATGTCAGGTCTCTGGT | 4284 | NEK6_6 | NEK6 | #65a2ae |
| GCATTCACGCCGGGTGATGCA | 15151 | NEK6_7 | NEK6 | #65a2ae |
| AGGCGAGGCAGGACTGTGTCA | 19174 | NEK6_8 | NEK6 | #65a2ae |
| GCTACGACTATGAGTTCGACC | 3393 | AKAP8_1 | AKAP8 | #65a2ae |
| CGTGGACCACAATCACAACCG | 16287 | AKAP8_2 | AKAP8 | #65a2ae |
| CCCTGGAACGAGCTGAACTAC | 68123 | AKAP8_3 | AKAP8 | #65a2ae |
| CAGCTTGGTGCTTATGAACCG | 9106 | AKAP8_4 | AKAP8 | #65a2ae |
| AGTCTAAAGACGCTGTTCCCA | 22060 | AKAP8_5 | AKAP8 | #65a2ae |
| GATCAACAGTTTCACTGGTGA | 1467 | AKAP8_6 | AKAP8 | #65a2ae |
| CGCAGCTGGTGACTTCCGCTC | 35781 | AKAP8_7 | AKAP8 | #65a2ae |
| GCTATGACAGCACCATGCCCT | 47764 | AKAP8_8 | AKAP8 | #65a2ae |
| AGCATCCCCGTCGATTGCATG | 12836 | DCHS1_1 | DCHS1 | #65a2ae |
| CACGTCGGGGTCGGTTGCTCG | 22615 | DCHS1_2 | DCHS1 | #65a2ae |
| TACTACAAGCTGATGTCGAGC | 5982 | DCHS1_3 | DCHS1 | #65a2ae |
| AGGTGACTCGACTGTTAACAC | 8317 | DCHS1_4 | DCHS1 | #65a2ae |
| GCCACTGGACTTTGAGCAGCG | 28116 | DCHS1_5 | DCHS1 | #65a2ae |
| TGCCACCGTAGAAGTTACAGT | 27897 | DCHS1_6 | DCHS1 | #65a2ae |
| AGGTCAGACGTCCATTGGGTC | 21350 | DCHS1_7 | DCHS1 | #65a2ae |
| CCCCGACTCAGGTGTTAACAG | 27349 | DCHS1_8 | DCHS1 | #65a2ae |
| TCTGGTGGCCGAAGCCGTAGT | 6644 | HTT_1 | HTT | #65a2ae |
| GGTCGACATCCTTGCTTGTCG | 15679 | HTT_2 | HTT | #65a2ae |
| TGTGACGCTGACAGAACTGCG | 14960 | HTT_3 | HTT | #65a2ae |
| CTGTAGTCCACATACCCATGG | 1379 | HTT_4 | HTT | #65a2ae |
| TGGGAACCAAGCTGACGAGAG | 23193 | HTT_5 | HTT | #65a2ae |
| CAGAGGGGCTCATCATTATAC | 18836 | HTT_6 | HTT | #65a2ae |
| TGCACGGCGTCCTCTATGTGC | 8566 | HTT_7 | HTT | #65a2ae |
| CATTTCTACCCGGCGACAAGC | 33222 | HTT_8 | HTT | #65a2ae |
| GATGTACATACCTTGCTTTGG | 29125 | BBIP1_1 | BBIP1 | #65a2ae |
| GAGAAAATGCATCAAGCAGCA | 12241 | BBIP1_2 | BBIP1 | #65a2ae |
| TAAACTTAGGGCCACTGTTTG | 18165 | BBIP1_3 | BBIP1 | #65a2ae |
| TATCCAACAACTCAGATATGG | 17238 | BBIP1_4 | BBIP1 | #65a2ae |
| TGTGATGTACATACCTTGCTT | 24651 | BBIP1_5 | BBIP1 | #65a2ae |
| AACAACTCAGATATGGCAGAA | 71381 | BBIP1_6 | BBIP1 | #65a2ae |
| TTGTGATGTACATACCTTGCT | 26381 | BBIP1_7 | BBIP1 | #65a2ae |
| CCTTGCTTTGGAAGAACTTCC | 35826 | BBIP1_8 | BBIP1 | #65a2ae |

### Table S2.

| | |
|---|---|
| CGGAAAACGCTCGACA | 0 |
| GGAAAACGCTCGACAA | 0 |
| TCGGACGCTGTCACGG | 0 |
| AAGCTCAGCCTGGCTG | 0 |
| ATAAAATTGGCACGGA | 0 |
| TGAAGGAAATAAAATT | 0 |
| GCCTGAGCCCGTCTAC | 0 |
| GAGGTAGACGGGCTCA | 0 |
| AACCACTTGAAAGATC | 0 |
| TCCCCTAGCGATCTTT | 0 |
| CAGCGGAGATTCTCCG | 0 |
| CTTCTGCCTGAGAATT | 0 |
| CCGCTTGGGTCCCAGG | 0 |
| GCGCCGCGCTCGAGGC | 0 |
| TCAACAGTAACTTCCG | 0 |
| TTGTCACTCTACCCTG | 0 |
| GTCTACGCAGGTCCGG | 0 |
| GGCCAGGACAGCCAAC | 0 |
| TTGCCCGGCCTACGCA | 0 |
| GCGTAGGCCGGGCAAC | 0 |
| GGACCCGAGCGCGAAG | 0 |
| TCCCGCTTGTTGGGGG | 0 |
| TGCCGCCACGCAAGCT | 0 |
| CGCCCCTCAAGCTTGC | 0 |
| CCATTGGCTGGTCTTC | 0 |
| CTGGATCGTAGCAGCC | 0 |
| GGGGCGGCGGCGCGGA | 0 |
| AGGGGCGGAGGGAGGC | 0 |
| TTGGCAGAGTCCGCAG | 0 |
| GTCAATCATGCTGGCC | 0 |
| CCTCACTAGTCAAGCC | 0 |
| GCCTCAATAACCCTAA | 0 |
| CCTTTTCTTGATTGTC | 0 |
| CCTCTGCCGCCCCTTTC | 0 |
| CGTGCATGTCTGTTCA | 0 |
| GACGATCAATTCAAGA | 0 |
| ATTCTGTACGACACTT | 0 |
| CCAGTGATGCACAATC | 0 |
| TGTCTCCGCACGCCGAAGTGA | 70566 |
| AAGATCCTAGAACAGATAGAC | 52362 |
| ACCAAAGCAAAACGATCTACC | 48062 |
| AGAGTGTTATATGTGTCATCG | 54421 |
| GCAGCTTATAATCTAGCACCG | 50822 |
| ATTATTAGACCATTCCTCCAG | 61386 |
| AGTCAGCTTCAAAGACGTGGC | 96451 |
| CTCATCGATGTAGACGATGCA | 33456 |
| TTCACATACGACTGACGGCCA | 66286 |
| CAGGCTGAAAAAACAGAACGA | 47746 |
| TCAAGTGCGATGCCATATAAG | 47399 |
| GTGAAAGCAGAATTATGTCTG | 47410 |
| CTGTTGGCTCGATTGAAACTT | 38938 |
| GAAGCCTGCTTTCTCTCGAGG | 59033 |
| TCTGGTTAGGCACTCCAACAG | 49652 |
| GTAAACTTGACGGACTAATAA | 45714 |
| GAAAAGTGAGGTACATGACCT | 41676 |
| CTTCTTTAGTCGCACTACGTA | 59416 |
| AATCCAGATCGCATCAAACAA | 75751 |
| GGATACAGATCTCCAAAAAAG | 34078 |
| ATCCGATGTCAGGTCTCTGGT | 42471 |
| AGGCGAGGCAGGACTGTGTCA | 49545 |
| CAGCTTGGTGCTTATGAACCG | 20176 |
| GATCAACAGTTTCACTGGTGA | 66672 |
| AGGTGACTCGACTGTTAACAC | 42586 |
| AGGTCAGACGTCCATTGGGTC | 57177 |
| GGTCGACATCCTTGCTTGTCG | 48914 |
| CTGTAGTCCACATACCCATGG | 12978 |
| GAGAAAATGCATCAAGCAGCA | 45262 |
| TGTGATGTACATACCTTGCTT | 39318 |
| TGTTTAAGGCCTATGTTTACGT | 47852 |
| TAGGCCACCGGATAAGGATCTT | 28372 |
| AGAAGCTACCGCGAGTTGAGTC | 54625 |
| ATAAAGCGGATCATTGCCTCGA | 61955 |
| TTAGGTCTTTGGGCCACAGGCG | 70363 |
| TCGAGGCAAATACTAATAGGGG | 27812 |
| ACAGTCACCACGATAACACAGC | 51087 |
| CGTCGGTACTTCACGGGTTAAG | 64322 |
| scriptcontrol | 0 |

| | |
|---|---|
| CGGAAAACGCTCGACA | 54697 |
| GGAAAACGCTCGACAA | 87757 |
| TCGGACGCTGTCACGG | 38274 |
| AAGCTCAGCCTGGCTG | 22448 |
| ATAAAATTGGCACGGA | 64793 |
| TGAAGGAAATAAAATT | 64369 |
| GCCTGAGCCCGTCTAC | 38693 |
| GAGGTAGACGGGCTCA | 25280 |
| AACCACTTGAAAGATC | 108190 |
| TCCCCTAGCGATCTTT | 94608 |
| CAGCGGAGATTCTCCG | 18182 |
| CTTCTGCCTGAGAATT | 91923 |
| CCGCTTGGGTCCCAGG | 25779 |
| GCGCCGCGCTCGAGGC | 9490 |
| TCAACAGTAACTTCCG | 28810 |
| TTGTCACTCTACCCTG | 45711 |
| GTCTACGCAGGTCCGG | 77085 |
| GGCCAGGACAGCCAAC | 21065 |
| TTGCCCGGCCTACGCA | 56638 |
| GCGTAGGCCGGGCAAC | 59899 |
| GGACCCGAGCGCGAAG | 26212 |
| TCCCGCTTGTTGGGGG | 69564 |
| TGCCGCCACGCAAGCT | 58567 |
| CGCCCCTCAAGCTTGC | 87448 |
| CCATTGGCTGGTCTTC | 27974 |
| CTGGATCGTAGCAGCC | 113624 |
| GGGGCGGCGGCGCGGA | 19950 |
| AGGGGCGGAGGGAGGC | 39879 |
| TTGGCAGAGTCCGCAG | 71108 |
| GTCAATCATGCTGGCC | 29234 |
| CCTCACTAGTCAAGCC | 77156 |
| GCCTCAATAACCCTAA | 16643 |
| CCTTTTCTTGATTGTC | 50904 |
| CCTCTGCCGCCCCTTTC | 45325 |
| CGTGCATGTCTGTTCA | 85550 |
| GACGATCAATTCAAGA | 105591 |
| ATTCTGTACGACACTT | 82766 |
| CCAGTGATGCACAATC | 43570 |
| TGTCTCCGCACGCCGAAGTGA | 0 |
| AAGATCCTAGAACAGATAGAC | 0 |
| ACCAAAGCAAAACGATCTACC | 0 |
| AGAGTGTTATATGTGTCATCG | 0 |
| GCAGCTTATAATCTAGCACCG | 0 |
| ATTATTAGACCATTCCTCCAG | 0 |
| AGTCAGCTTCAAAGACGTGGC | 0 |
| CTCATCGATGTAGACGATGCA | 0 |
| TTCACATACGACTGACGGCCA | 0 |
| CAGGCTGAAAAAACAGAACGA | 0 |
| TCAAGTGCGATGCCATATAAG | 0 |
| GTGAAAGCAGAATTATGTCTG | 0 |
| CTGTTGGCTCGATTGAAACTT | 0 |
| GAAGCCTGCTTTCTCTCGAGG | 0 |
| TCTGGTTAGGCACTCCAACAG | 0 |
| GTAAACTTGACGGACTAATAA | 0 |
| GAAAAGTGAGGTACATGACCT | 0 |
| CTTCTTTAGTCGCACTACGTA | 0 |
| AATCCAGATCGCATCAAACAA | 0 |
| GGATACAGATCTCCAAAAAAG | 0 |
| ATCCGATGTCAGGTCTCTGGT | 0 |
| AGGCGAGGCAGGACTGTGTCA | 0 |
| CAGCTTGGTGCTTATGAACCG | 0 |
| GATCAACAGTTTCACTGGTGA | 0 |
| AGGTGACTCGACTGTTAACAC | 0 |
| AGGTCAGACGTCCATTGGGTC | 0 |
| GGTCGACATCCTTGCTTGTCG | 0 |
| CTGTAGTCCACATACCCATGG | 0 |
| GAGAAAATGCATCAAGCAGCA | 0 |
| TGTGATGTACATACCTTGCTT | 0 |
| TGTTTAAGGCCTATGTTTACGT | 0 |
| TAGGCCACCGGATAAGGATCTT | 0 |
| AGAAGCTACCGCGAGTTGAGTC | 0 |
| ATAAAGCGGATCATTGCCTCGA | 0 |
| TTAGGTCTTTGGGCCACAGGCG | 0 |
| TCGAGGCAAATACTAATAGGGG | 0 |
| ACAGTCACCACGATAACACAGC | 0 |
| CGTCGGTACTTCACGGGTTAAG | 0 |
| scriptcontrol | 0 |

| | |
|---|---|
| CGGAAAACGCTCGACAACGG | 170620 |
| GGAAAACGCTCGACAACGGA | 254924 |
| TCGGACGCTGTCACGGCTCG | 138695 |
| AAGCTCAGCCTGGCTGCGCA | 58979 |
| ATAAAATTGGCACGGAGTCT | 195978 |
| TGAAGGAAATAAAATTGGCA | 177181 |
| GCCTGAGCCCGTCTACCTCG | 324869 |
| GAGGTAGACGGGCTCAGGCT | 70045 |
| AACCACTTGAAAGATCGCTA | 430280 |
| TCCCCTAGCGATCTTTCAAG | 313579 |
| CAGCGGAGATTCTCCGCAGC | 75837 |
| CTTCTGCCTGAGAATTGACT | 394551 |
| CCGCTTGGGTCCCAGGGCCG | 68716 |
| GCGCCGCGCTCGAGGCGGCC | 31290 |
| TCAACAGTAACTTCCGATCG | 92851 |
| TTGTCACTCTACCCTGCTGG | 107876 |
| GTCTACGCAGGTCCGGAAGT | 338482 |
| GGCCAGGACAGCCAACCTGC | 54199 |
| TTGCCCGGCCTACGCATCGG | 165951 |
| GCGTAGGCCGGGCAACATGG | 159393 |
| GGACCCGAGCGCGAAGTGTG | 72315 |
| TCCCGCTTGTTGGGGGGGGC | 163408 |
| TGCCGCCACGCAAGCTTGAG | 172842 |
| CGCCCCTCAAGCTTGCGTGG | 311986 |
| CCATTGGCTGGTCTTCACGA | 65959 |
| CTGGATCGTAGCAGCCCGAA | 427478 |
| GGGGCGGCGGCGCGGAGCGC | 53808 |
| AGGGGCGGAGGGAGGCCGGG | 130720 |
| TTGGCAGAGTCCGCAGGCTA | 286209 |
| GTCAATCATGCTGGCCGGCG | 78181 |
| CCTCACTAGTCAAGCCGAGA | 316073 |
| GCCTCAATAACCCTAACAGC | 41010 |
| CCTTTTCTTGATTGTCGAGT | 149232 |
| CCTCTGCCGCCCCTTTCACAC | 139533 |
| CGTGCATGTCTGTTCATAAC | 226057 |
| GACGATCAATTCAAGATGTA | 381688 |
| ATTCTGTACGACACTTTAGT | 299607 |
| CCAGTGATGCACAATCTATC | 166623 |
| TGTCTCCGCACGCCGAAGTGA | 0 |
| AAGATCCTAGAACAGATAGAC | 0 |
| ACCAAAGCAAAACGATCTACC | 0 |
| AGAGTGTTATATGTGTCATCG | 0 |
| GCAGCTTATAATCTAGCACCG | 0 |
| ATTATTAGACCATTCCTCCAG | 0 |
| AGTCAGCTTCAAAGACGTGGC | 0 |
| CTCATCGATGTAGACGATGCA | 0 |
| TTCACATACGACTGACGGCCA | 0 |
| CAGGCTGAAAAAACAGAACGA | 0 |
| TCAAGTGCGATGCCATATAAG | 0 |
| GTGAAAGCAGAATTATGTCTG | 0 |
| CTGTTGGCTCGATTGAAACTT | 0 |
| GAAGCCTGCTTTCTCTCGAGG | 0 |
| TCTGGTTAGGCACTCCAACAG | 0 |
| GTAAACTTGACGGACTAATAA | 0 |
| GAAAAGTGAGGTACATGACCT | 0 |
| CTTCTTTAGTCGCACTACGTA | 0 |
| AATCCAGATCGCATCAAACAA | 0 |
| GGATACAGATCTCCAAAAAAG | 0 |
| ATCCGATGTCAGGTCTCTGGT | 0 |
| AGGCGAGGCAGGACTGTGTCA | 0 |
| CAGCTTGGTGCTTATGAACCG | 0 |
| GATCAACAGTTTCACTGGTGA | 0 |
| AGGTGACTCGACTGTTAACAC | 0 |
| AGGTCAGACGTCCATTGGGTC | 0 |
| GGTCGACATCCTTGCTTGTCG | 0 |
| CTGTAGTCCACATACCCATGG | 0 |
| GAGAAAATGCATCAAGCAGCA | 0 |
| TGTGATGTACATACCTTGCTT | 0 |
| CGTCGGTACTTCACGGGTTAAG | 0 |
| TGTTTAAGGCCTATGTTTACGT | 0 |
| TAGGCCACCGGATAAGGATCTT | 0 |
| AGAAGCTACCGCGAGTTGAGTC | 0 |
| ATAAAGCGGATCATTGCCTCGA | 0 |
| TTAGGTCTTTGGGCCACAGGCG | 0 |
| TCGAGGCAAATACTAATAGGGG | 0 |
| ACAGTCACCACGATAACACAGC | 0 |

| | |
|---|---|
| CGGAAAACGCTCGACAACGG | 0 |
| GGAAAACGCTCGACAACGGA | 0 |
| TCGGACGCTGTCACGGCTCG | 0 |
| AAGCTCAGCCTGGCTGCGCA | 0 |
| ATAAAATTGGCACGGAGTCT | 0 |
| TGAAGGAAATAAAATTGGCA | 0 |
| GCCTGAGCCCGTCTACCTCG | 0 |
| GAGGTAGACGGGCTCAGGCT | 0 |
| AACCACTTGAAAGATCGCTA | 0 |
| TCCCCTAGCGATCTTTCAAG | 0 |
| CAGCGGAGATTCTCCGCAGC | 0 |
| CTTCTGCCTGAGAATTGACT | 0 |
| CCGCTTGGGTCCCAGGGCCG | 0 |
| GCGCCGCGCTCGAGGCGGCC | 0 |
| TCAACAGTAACTTCCGATCG | 0 |
| TTGTCACTCTACCCTGCTGG | 0 |
| GTCTACGCAGGTCCGGAAGT | 0 |
| GGCCAGGACAGCCAACCTGC | 0 |
| TTGCCCGGCCTACGCATCGG | 0 |
| GCGTAGGCCGGGCAACATGG | 0 |
| GGACCCGAGCGCGAAGTGTG | 0 |
| TCCCGCTTGTTGGGGGGGGC | 0 |
| TGCCGCCACGCAAGCTTGAG | 0 |
| CGCCCCTCAAGCTTGCGTGG | 0 |
| CCATTGGCTGGTCTTCACGA | 0 |
| CTGGATCGTAGCAGCCCGAA | 0 |
| GGGGCGGCGGCGCGGAGCGC | 0 |
| AGGGGCGGAGGGAGGCCGGG | 0 |
| TTGGCAGAGTCCGCAGGCTA | 0 |
| GTCAATCATGCTGGCCGGCG | 0 |
| CCTCACTAGTCAAGCCGAGA | 0 |
| GCCTCAATAACCCTAACAGC | 0 |
| CCTTTTCTTGATTGTCGAGT | 0 |
| CCTCTGCCGCCCCTTTCACAC | 0 |
| CGTGCATGTCTGTTCATAAC | 0 |
| GACGATCAATTCAAGATGTA | 0 |
| ATTCTGTACGACACTTTAGT | 0 |
| CCAGTGATGCACAATCTATC | 0 |
| TGTCTCCGCACGCCGAAGTGA | 231335 |
| AAGATCCTAGAACAGATAGAC | 195306 |
| ACCAAAGCAAAACGATCTACC | 161495 |
| AGAGTGTTATATGTGTCATCG | 188679 |
| GCAGCTTATAATCTAGCACCG | 132178 |
| ATTATTAGACCATTCCTCCAG | 234354 |
| AGTCAGCTTCAAAGACGTGGC | 324429 |
| CTCATCGATGTAGACGATGCA | 194094 |
| TTCACATACGACTGACGGCCA | 178060 |
| CAGGCTGAAAAAACAGAACGA | 186883 |
| TCAAGTGCGATGCCATATAAG | 160762 |
| GTGAAAGCAGAATTATGTCTG | 158399 |
| CTGTTGGCTCGATTGAAACTT | 127424 |
| GAAGCCTGCTTTCTCTCGAGG | 349266 |
| TCTGGTTAGGCACTCCAACAG | 169960 |
| GTAAACTTGACGGACTAATAA | 189205 |
| GAAAAGTGAGGTACATGACCT | 110015 |
| CTTCTTTAGTCGCACTACGTA | 272311 |
| AATCCAGATCGCATCAAACAA | 284963 |
| GGATACAGATCTCCAAAAAAG | 220720 |
| ATCCGATGTCAGGTCTCTGGT | 142862 |
| AGGCGAGGCAGGACTGTGTCA | 170252 |
| CAGCTTGGTGCTTATGAACCG | 115283 |
| GATCAACAGTTTCACTGGTGA | 190982 |
| AGGTGACTCGACTGTTAACAC | 142684 |
| AGGTCAGACGTCCATTGGGTC | 240633 |
| GGTCGACATCCTTGCTTGTCG | 230646 |
| CTGTAGTCCACATACCCATGG | 35182 |
| GAGAAAATGCATCAAGCAGCA | 165051 |
| TGTGATGTACATACCTTGCTT | 201646 |
| CGTCGGTACTTCACGGGTTAAG | 271823 |
| TGTTTAAGGCCTATGTTTACGT | 171236 |
| TAGGCCACCGGATAAGGATCTT | 91287 |
| AGAAGCTACCGCGAGTTGAGTC | 152365 |
| ATAAAGCGGATCATTGCCTCGA | 219112 |
| TTAGGTCTTTGGGCCACAGGCG | 292320 |
| TCGAGGCAAATACTAATAGGGG | 135801 |
| ACAGTCACCACGATAACACAGC | 149646 |

**Supplementary Table 2**: Genomic and Plasmid DNA raw read counts from mosaic double perturbed organoids (from left to right: saCas9 plasmid; spCas9 plasmid; spCas9 genomic ; saCas9 genomic
